# Unilateral loss of *recql4* function in *Xenopus laevis* tadpoles leads to ipsilateral ablation of the forelimb, hypoplastic Meckel’s cartilage and vascular defects

**DOI:** 10.1101/2025.04.14.648861

**Authors:** Caroline W. Beck, Matthew Reily-Bell, Louise S. Bicknell

**Author notes:** Correspondence to: Associate Professor Caroline Beck.

## Abstract

*RECQL4* encodes a RecQ helicase, one of a family of DNA unwinding enzymes with roles in DNA replication, double strand break repair and genomic stability. Pathogenic variants in *RECQL4* are clinically associated with three rare autosomal recessive conditions: Rothmund-Thomson Syndrome type II, Baller-Gerold Syndrome and RAPADILINO syndrome. These three syndromes show overlapping growth retardation, low bone density and skeletal defects affecting the arms and hands. Here, we take advantage of the ability to generate one-sided CRISPR knockdowns of *recql4* in *Xenopus laevis* tadpoles. Tadpoles develop normally until feeding starts, after which growth slows on the edited side leading to a curved posture, smaller eyes (micropthalmia) and reduced head size (microcephaly). Forelimb buds fail to develop, leading to complete absense of the forelimb on the edited side. Additionally, Meckel’s cartilage (lower jaw) ossification is absent or reduced and the hyoid cartilage is smaller, but this is not due to deficiencies in cranial neural crest migration on the edited side. Knockdown of *recql4* also results in hypoplastic vasculature, with reduced branching from the aorta on the edited side. Taken together, our results clearly show the utility of unilateral CRISPR editing in *Xenopus* for understanding the specific phenotypic developmental effects of mutations affecting cell proliferation.

## Introduction

Many developmental syndromes result from genetic alterations that disrupt timely and accurate DNA replication and DNA repair (Ichikawa *et al*., 2002, Hoki *et al*., 2003, Mann *et al*., 2005, Bohr, 2008, Siitonen *et al*., 2009, Larsen & Hickson, 2013, Croteau *et al*., 2014, Smeets *et al*., 2014, Lu *et al*., 2015, Kaiser *et al*., 2017, Castillo-Tandazo *et al*., 2019, Bellelli & Boulton, 2021, Luong & Bernstein, 2021). RECQL4 belongs to a family of DNA helicases and has been shown to be required for DNA replication in *Xenopus laevis egg extracts* (Sangrithi *et al*., 2005). Additionally, RECQL4 stabilises and repairs replication forks and plays a role in repairing double stranded DNA breaks, (Singh *et al*., 2010, Lu *et al*., 2017, Luong & Bernstein, 2021). In humans, biallelic loss of function variants of *RECQL4* have been associated with three rare genetic conditions (RECQL4 syndromes): Rothmund Thomson syndrome (RTS) (Kitao *et al*., 1999), Baller Gerald syndrome (BGS) (Van Maldergem *et al*., 2006), and RAPADILINO syndrome (Siitonen *et al*., 2003). While RAPADILINO syndrome mainly results from a single *RECQL4* variant more common in the Finnish population, BGS and RTS are genetically heterogeneous. All three RECQL4 syndromes show core overlapping phenotypes of growth retardation (leading to proportional short stature), radial ray defects in upper limbs (especially loss of thumbs), and osteopenia (low bone density) (Siitonen *et al*., 2009, Luong & Bernstein, 2021). Additional phenotypes common in RTS type II are poikiloderma, a skin disorder shared with Bloom syndrome, and predisposition to osteosarcoma (Kitao *et al*., 1999, Wang *et al*., 2003). More recently, variants have been associated with myelodysplastic syndrome in multiple generations of a single family, although no functional experiments to test the missense variants were undertaken (Jiang *et al*., 2023). Whether there are truly clinically distinct disorders caused by the disruption of the same gene rather than just very broad and heterogeneous clinical presentations is debated, (Van Maldergem *et al*., 2016, Martins *et al*., 2023) especially as the number of patients increase and genetic testing expands.

*RECQL4* is a member of the RecQ helicase gene family. While prokaryotes and yeast contain only one RecQ gene, in humans and *Xenopus*, there are five: *RECQL, BLM/RECQL2, WRN/RECQL3, RECQL4 and RECQL5*. The proteins produced by these genes all function in genome maintenance and stability and are often referred to as “guardians of the genome” (Hickson, 2003, Bohr, 2008, Larsen & Hickson, 2013, Croteau *et al*., 2014). In addition to *RECQL4*, developmental syndromes have been attributed to genetic impairment of *RECQL* (RECON syndrome), *WRN* (Werner syndrome) and *BLM* (Bloom syndrome) (Abu-Libdeh *et al*., 2022). The human RecQ helicases share some core domains such as the helicase, but are also divergent in structure, function, cellular localization and protein-protein interactions, suggesting division of labour (Croteau *et al*., 2014).

While sharing many phenotypic similarities such as cancer predisposition and premature aging, RECQL4-associated syndromes are unique among the RECQ helicase family for the strong presence of skeletal abnormalities (Arora *et al*., 2014, Oshima *et al*., 2017, Abu-Libdeh *et al*., 2022). More than 100 *RECQL4* pathogenic variants have been reported, with most either nonsense or frameshift variants predicted to results in either null alleles (due to premature termination codons triggering nonsense-mediated decay pathways) or truncated proteins that would disrupt the helicase domain (Siitonen *et al*., 2009, Castillo-Tandazo *et al*., 2021). The C-terminal region of RECQL4 (RQC) is also known to be essential for activity (Mojumdar *et al*., 2017) and the most common variant associated with BGS is in this region (Luong & Bernstein, 2021).

The exact mechanisms by which *RECQL4* variants cause the broad range of observed phenotypes are yet to be elucidated and several animal models have been developed. *Drosophila melanogaster* homozygous for null *recql4* mutations are lethal, and helicase-dead Recql4 does not complement this, suggesting that the helicase activity is critical for viability (Capp *et al*., 2009). Similarly, deletion of all or part of the helicase domain in mice results in embryonic lethality (Hoki *et al*., 2003, Smeets *et al*., 2014). However, deletion of exons 5-8, which covers part of the N-terminal SLD-2 like DNA replication initiation domain of Recql4, also causes early embryonic lethality, even though the helicase domain (exons 8-14) is intact (Ichikawa *et al*., 2002). A viable mouse model replicating the human skin and skeletal phenotypes was made by replacing exons 9-13 with a mini-gene cassette harboring termination codons, which created truncated *Recql4* transcripts (Mann *et al*., 2005). Additionally, a substitution designed to remove the ATPase required for helicase function (K525A) did not cause adverse developmental effects in mice (Castillo-Tandazo *et al*., 2019). There may be differences in expression of RECQL4 that explain these different outcomes in mice and humans. Conditional mouse *recql4* mutants affecting the limb bud mesenchyme (Prx1-Cre) were viable and replicated limb skeletal defects seen in patients (Lu *et al*., 2015).

In order to determine how loss of function of RECQL4 leads to specific developmental skeletal phenotypes in humans, such as short stature and limb defects, we turned to the model organism *Xenopus laevis*. *Xenopus* species are increasingly being used to model human genetic disease due to a high level of genetic conservation, ease of CRISPR/Cas9 editing, and well-characterized and observable development which occurs externally (reviewed in (Willsey *et al*., 2024)). While *X. laevis* is allotetraploid, with around 60% of genes having two homeologues, there is a single copy of *recql4*, on chromosome 2L (*recql4.L*, xenbase.org, (Nenni *et al*., 2019, Fisher *et al*., 2023)). Additionally, it is possible to generate embryos and tadpoles that are edited only on one side of the body, by targeting CRISPR reagents to one cell of the two-cell stage embryo (Willsey *et al*., 2021). Here, we take advantage of this to generate one-sided CRISPR/cas9 edited models of RECQL4 syndromes, in which the unedited side acts as an internal control for the phenotype. Human genetic variants commonly interrupt the helicase domain (Wang *et al*., 2003, Larizza *et al*., 2010, Martins *et al*., 2023), although the precise protein-level consequences have often not been experimentally determined. We therefore designed our gRNA to approximate human variants by disrupting the helicase domain. Edited tadpoles show ipsilateral phenotypes of reduced growth, absence of forelimb (and more rarely hindlimb) development, and specific craniofacial cartilage defects, all in keeping with, but more severe than, the phenotypes observed in humans. Intriguingly, we also observed hypoplastic aorta and failure to form the subclavian arterial branch on the edited side, which has not been reported previously. Our work shows the utility of *Xenopus* one-sided CRISPants to model disorders of growth caused by pathogenic variants in genes associated with replication, helping to explain how loss of function leads specific developmental phenotypes.

## Methods

### Animal ethics

*Xenopus laevis* adults were held under University of Otago Animal Ethics Committee animal use procedure AUP-22-24, and embryos were produced according to AUP-22-12. CRISPR modification with *recql4* sgRNAs was approved and carried out according to AUP-25-04.

### Production of one-sided recql4 CRISPants

Adult female *Xenopus laevis* were induced to lay eggs by injection of 500 IU Chorulon (HCG) per 75 g bodyweight, into the dorsal lymph sac. After 16 hours, females were transferred into 1 x Marc’s modified ringers (MMR: 10 mM NaCl, 0.1 mM MgSO4.6H2O, 0.2 mM CaCl2, 0.5 mM HEPES, 10 μM EDTA, pH 7.8), from which eggs were collected for *in vitro* fertilisation. Jelly coats were removed from embryos after 20 minutes by gentle swirling in 2% L-Cysteine (pH 7.9) and rinsing three times in 1 x MMR.

Unique sgRNA sequences targeting *X. laevis recql4.L* (RefSeq: NM_001095713) were designed using ChopChop v3 (Labun *et al*., 2019). InDelphi (Shen *et al*., 2018) was used to predict editing efficiency in genomic context and to select guides predicted to cause a dominant frameshift mutation (Figure 1, Table 1). CRISPRScan (Moreno-Mateos *et al*., 2015) was used to confirm that guides had no off targets (none with <3 mismatches overall, or identical seed). The EnGen sgRNA oligo designer v1.12.1 tool (NEB) was used to generate 55 bp oligonucleotides which were synthesized by Integrated DNA Technologies (IDT). An EnGen Cas9 sgRNA kit (NEB) was used to generate sgRNA by *in vitro* transcription. sgRNA were purified using a Monarch RNA purification kit (NEB) and eluted into 30 μL pyrogen free water before storing as 3 μL aliquots at -70°C at 600 ng/μl. sgRNA were loaded onto Cas9-NLS (EnGen, *S. pyogenes*, NEB) as described in (Chapman *et al*., 2022). Synthetic capped mRNA for nGFP was prepared using an SP6 mMessage mMachine kit (Invitrogen) and diluted to 200 ng/μl to label the injection side. The injection mix was 0.3μl Cas9-NLS, 0.5 μl nGFP mRNA and 1 to 1.5 μl of sgRNA, made to 3 μl with nuclease free water. Two-cell stage embryos were transferred into 5% Ficoll 400 in 1 x MMR and placed in a well cut into a 2% agar lined 50 mm petri dish for injection. Cas9/sgRNA plus nGFP mRNA, or just the mRNA, were backloaded into a glass capillary needle (Drummond) pulled to a fine point using a Sutter P92 needle puller and filled with light paraffin oil. 10 nl was injected into one cell using a Nanoject III micropipette (Drummond) held with an MM3 micromanipulator. After 2-3 hours, embryos were transferred into 2.5% Ficoll in 0.1 x MMR and incubated at 23°C overnight.

**Figure 1.**
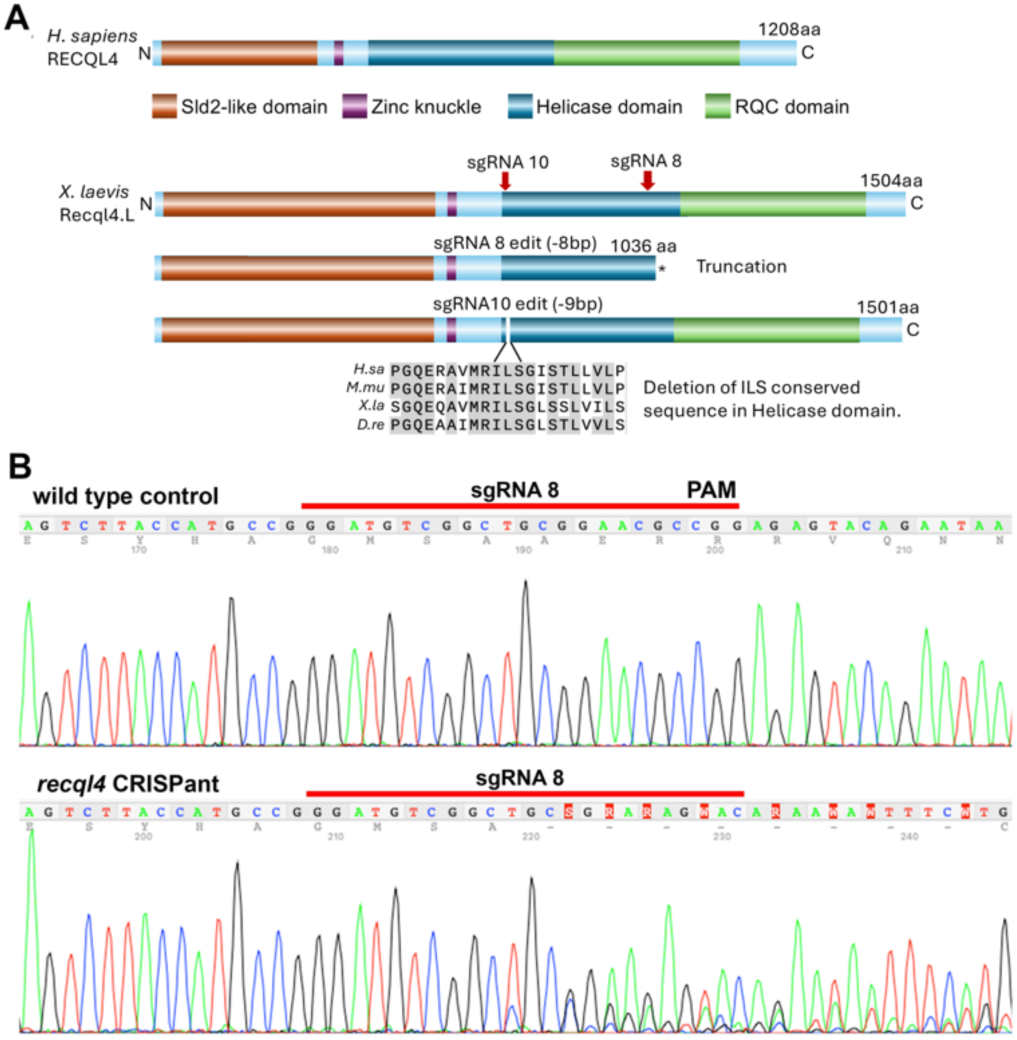
Schematic of *H. sapiens* and *X. laevis* RECQL4 proteins and CRISPR sgRNAs. **A)** Protein domain structure for human RECQL4 and *X. laevis* Recql4, red arrows indicate sgRNA target positions. Asterisk indicates premature stop codon. sgRNA 8 generates predominantly 8 bp deletions, that will introduce a downstream stop codon in the helicase domain, resulting in a truncated protein. sgRNA 10 generates a dominant 9 bp deletion that removes bases “ILS” in a conserved region of the helicase. Protein alignments are shown for human, mouse, *Xenopus* and zebrafish with grey boxes highlighting conserved residues. **B)** Example of sgRNA8 editing, Sanger sequence traces from control wild type and sgRNA8 CRISPant. Guide sequence is indicated by red line.

**Table 1.**
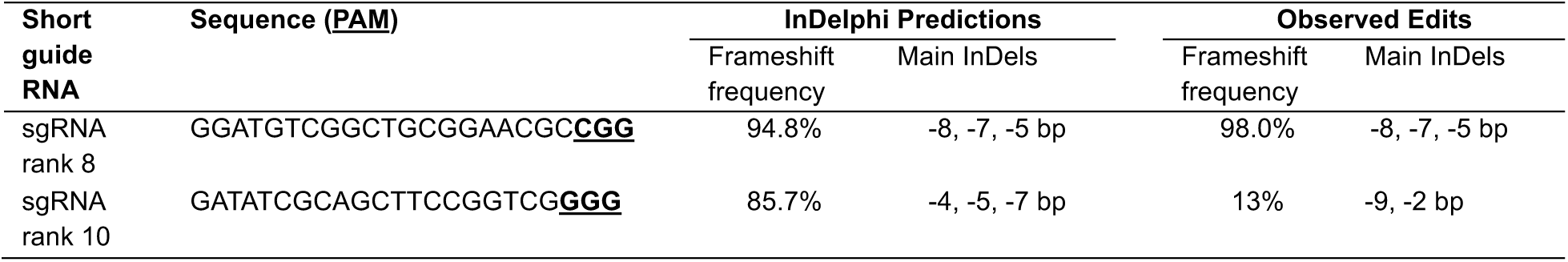
sgRNA, predicted and observed editing.

**Table 2.**
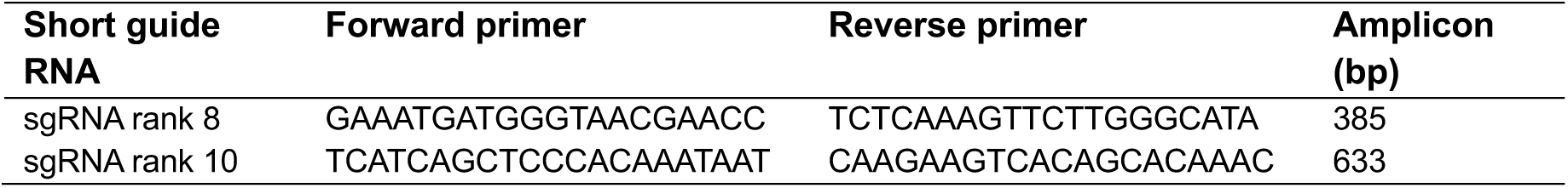
Primers for amplification and sequencing.

**Table 3.**
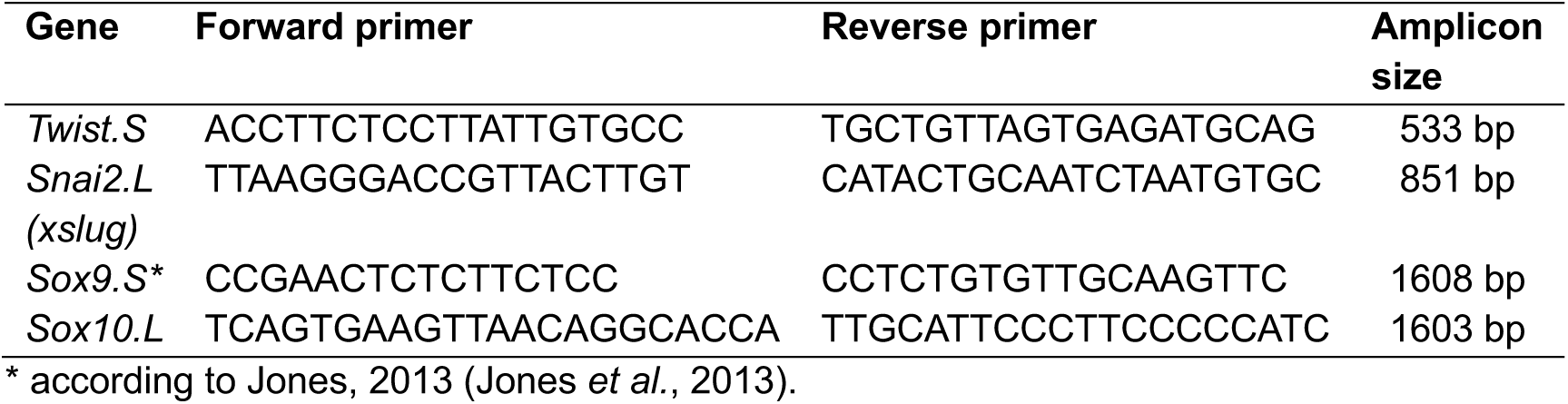
Primers used to generate ISH antisense probes.

Stage 25 neurula embryos were transferred into 0.1 x MMR and examined under epifluorescence to determine the location of the injection. Embryos with clearly left or right sided GFP were sorted and retained for phenotyping.

### Genotyping and editing analysis of CRISPants

For cohort genotyping, embryos with half green staining not tidily localised on the left or right side were randomly selected and collected into 0.2 μL PCR tube strips. Any liquid was removed and replaced with 150 μL of 5% Chelex 100 beads (Bio-Rad) suspended in Tris/EDTA buffer, pH 8.0. Each sample was homogenised by pipetting and 30 μg of Proteinase K (Sigma) was mixed in before incubating at 65 °C for 4 hours. Proteinase K was then inactivated by heating for 15 minutes at 95 °C. PCR primers around the editing site were designed using ChopChop v3 (Labun *et al*., 2019) (Table) and 1 μL of extracted DNA was PCR amplified for 35 cycles with MiFi mix (Bioline). A T7 endonuclease assay to detect heteroduplexes was used to confirm editing of around 50% in pooled amplicon samples, as expected for hemiCRISPants. Representative sample amplicons were cleaned using ExoSapIT (Applied Biosystems) and Sanger sequenced (Genetics Analysis Service, University of Otago). Sequence traces were deconvoluted by comparing to control unedited samples using TIDE v2 (Tracking of InDels by Decomposition) to confirm indel size (Brinkman *et al*., 2014).

### Phenotyping of unilateral CRISPants

Tadpoles were raised at 23°C in 10 mm petri dishes containing 0.1 x MMR until NF stage 47 (Nieuwkoop & Faber, 1994). They were then fed with 100 μL of fresh spirulina slurry and allowed to clear the dish overnight before moving to 1 L tanks and daily spirulina feeding. At stage 48/49 they were moved to tanks in a recirculating aquarium (Marine Biotech XR1) containing carbon filtered tap water, at a density of 25 tadpoles per litre. Tadpoles were fed daily with a slurry of spirulina powder with a small amount of ground adult frog pellets and recirculation turned off for two hours after each feed.

Tadpoles were anaesthetised twice briefly using 1/4000 v/v MS222 (Sigma) to observe development and take photographs. At stage 50 (approx. two weeks) the presence or absence of each limb bud was noted. At three weeks of age, 10 tadpoles per group were individually photographed on agar-lined plates using a Leica MZ FLIII stereomicroscope fitted with a Leica DFC320 camera and software. They were illuminated from underneath with an A4 LED light box (Huion) and from the sides using a swan neck light. FiJi image analysis (Schindelin *et al*., 2012) was used to obtain a quantification of left and right eye size and head size. Tadpoles were then recovered in fresh tanks with gentle air bubbling for 1 hour and returned to the aquarium. Every 3 days, tadpoles that had reached stage NF 58, defined by the hatching of the forelimbs from their pockets, were euthanised in 0.1% buffered benzocaine (Sigma), rinsed with 0.1 x MMR and fixed in 4% formaldehyde in phosphate buffered saline (PBS, Oxoid). Following fixation overnight, tails were removed and photographs taken of each tadpole.

Bone and cartilage staining followed the method of (Newman & Dumont, 1983) with modifications as in (Jones *et al*., 2013). Briefly, fixed tadpoles were eviscerated in PBS and bleached to remove skin pigmentation using 6% H2O2 in PBS. Alcian blue (0.01% in 40% acetic acid and 60% ethanol) was used to stain cartilage overnight, then destained in 70% ethanol. Tadpoles were then soaked in 1% KOH to clear the muscle tissues before staining the bone with alizarin red S (0.01% w/v in 1% KOH). Tadpoles were destained in 1% KOH and then mounted in 50% glycerol/50% ethanol for photography. Adobe Photoshop 2025 was used to prepare figures.

### In situ hybridisation

Whole mount colorimetric *in situ* hybridisation of stage 19 *recql4* half CRISPants was carried out as in Beck et al 1998 (Beck & Slack, 1998). Digoxygenin-UTP labelled antisense probes were visualised using anti-DIG Fab fragments conjugated to alkaline phosphatase followed by enzymatic conversion of NBT/BCIP (Invitrogen) to generate a dark blue precipitate. Embryos were bleached with 5% H2O2 post fixation.

### Graphing and statistical analysis

Data were graphed using GraphPad Prism v10.4.0. Quantitative data were first tested for normal distribution using the Shapiro-Wilk test, with appropriate selection of non-parametric tests for non-normal data. Categorical data were analysed using Fisher’s exact test.

## Results

### Unilateral knockdown of *recql4* results in smaller head and eye size on the edited side

Injection of cas9, 2-3 ng of *recql4* sgRNA8 or 10, and mRNA for nGFP into one cell at the 2-cell stage resulted in efficient editing for both sgRNAs in samples arbitrarily chosen 18 hours after injection (Figure 1B). The one cell injections resulted in a sharp left-right expression boundary of GFP in 60-70% of cases, clearly seen at neurula stages (Figure 2A, A’). We would therefore predict editing to be around 50% in these half-CRISPant embryos. A minimum of five individual neurula stage embryos for each guide were sequenced to determine the extent and type of editing. sgRNA8 gave high levels of frameshift edits as predicted by InDelphi, which would result in a premature stop codon at amino acid 1036, disrupting the helicase and removing the RQC domain (Figure 1, Table 1). Overall editing averaged 43.8% and 98% of edits resulted in frame shift. sgRNA10 caused predominantly 9 bp deletions which result in the loss of three amino acids in a highly conserved region of the helicase, and a 2 bp deletion. Frame shift levels averaged 13%, lower than predicted, with total editing efficiency averaging 30.0% (Figure 1, Table 1).

**Figure 2:**
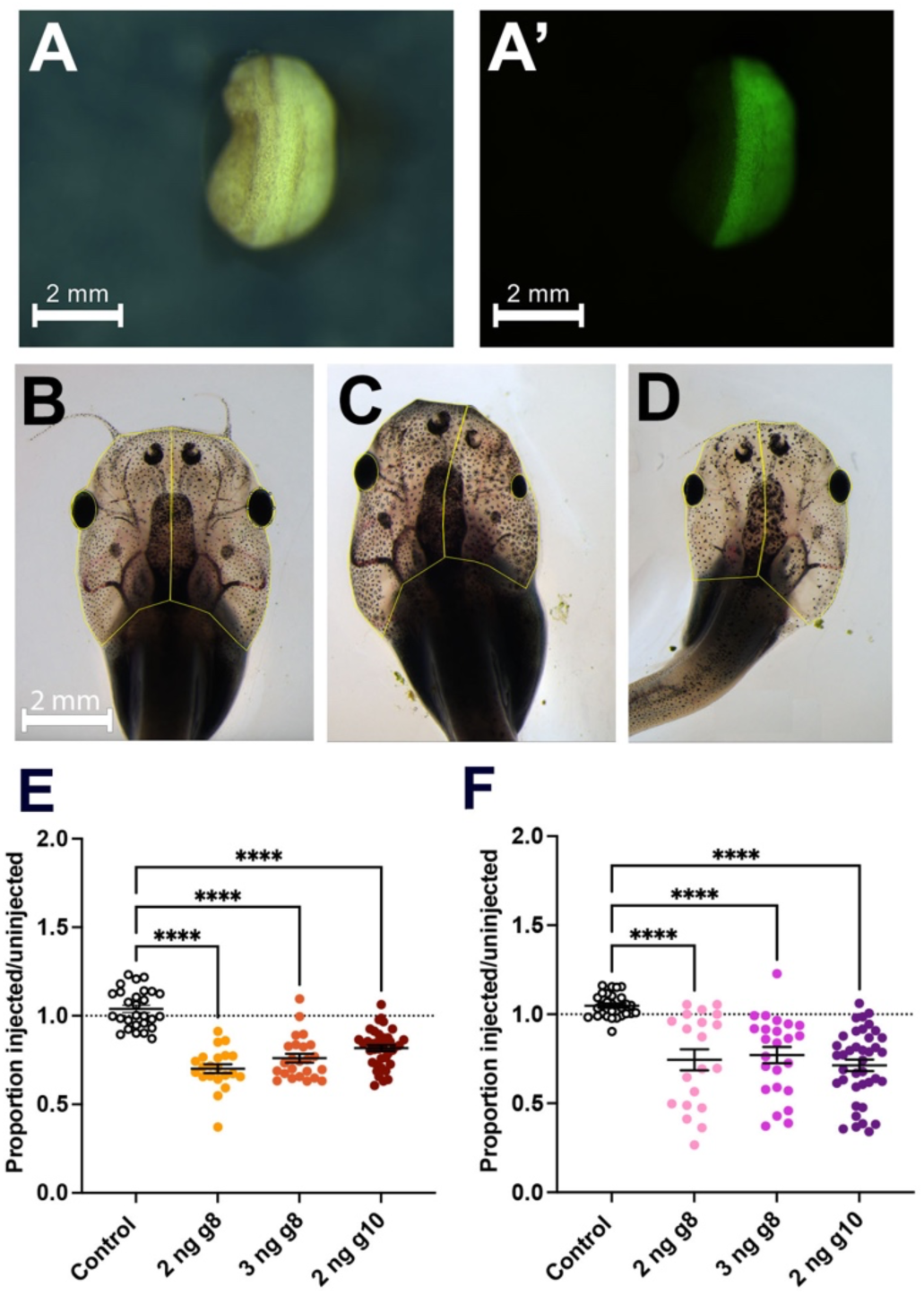
Unilateral knockdown of *recql4* results in smaller head and eye size on the edited side. **A)** Example of bright field and **A’**) fluorescence view of neurula embryo, dorsal view, anterior uppermost. Embryo was injected on the right side at the two-cell stage, GFP can be seen only on the right side of the midline. **B-D)** examples of tadpoles at 3 weeks, yellow outlines show the regions of interest (ROI) used to measure area of head and eye on left and right sides using FiJi. Scale bar in B applies to all panels. **B)** Control, injected with GFP mRNA only on the left side. **C)** Tadpole from the group injected with 2 ng *recql4* sgRNA 8 (g8) and nGFP mRNA on right, and **D)** on left. **E, F)** Scatter plots showing the proportions injected vs. uninjected head side (E) or eye (F) area. Line indicates mean ± SEM. Dotted line denotes no difference between the two sides. *Recql4* injected groups are compared to the control (only injected with GFP mRNA) using Kruskal-Wallis tests with Dunn’s post-hoc comparisons. Sample sizes are: GFP only controls: N=28, 2 ng g8 N=21, 3 ng g8 N=24, 2ng g10 N=39. Significantly different groups are indicated as p<0.0001****. Raw data can be found in Tables S1-S6 in File S1.

Using the co-injected transient GFP expression, embryos were sorted into clear left and clear right-sided cohorts and raised to tadpole stages. No obvious differences were observed at 5 days of age when feeding commenced. By three weeks of development, tadpoles had reached stage 50-52. Differences in the head and eye size on the side with *recql4* CRISPR knockdown with both sgRNA were clear in most individuals (Figure 2 B-D; Figure S1; Table S1, S2). Head size, estimated by comparing the area either side of the midline from dorsal view photographs, was 70% (+/- 2.0%) and 76% (+/- 2.5%) of the control side for sgRNA 8 at 2 and 3 ng doses and 82% (+/- 2.0%) of the control side for sgRNA10 on the uninjected side of one-sided CRISPants (Figure 2). Control tadpoles (only injected with GFP) showed no difference between left and right sides and were closest in size to the wild type halves. Taken together, this data indicates that *Recql4* knockdown limits growth of the tadpole head to 70-80% of normal size (Figure 2E).

Similarly, eye size (estimated as the area of an ellipse covering the retina from the same photographs) was often noted to be smaller on the CRISPant side (Figure 2, Figure S1). Eye area on the injected side was on average 74% (+/-6.0%) and 77% (+/- 4.5%) of control eye area for sgRNA8 at 2 and 3 ng respectively) and 71% (+/- 3.2%) for sgRNA10). Knockdown of *recql4* therefore also results in smaller size eyes, or micropthalmia (Figure 2F).

### Forelimb development requires Recql4

NF58 tadpoles from the sgRNA 8 cohorts were pooled and examined for cartilage and skeletal defects. At stage NF58, the forelimbs have been released from the flap of skin under which they develop and can be observed by eye for the first time (Figure 3, Figure S2, S3). The most frequently observed phenotype was a complete absence of the forelimb on the side of the tadpole with *recql4* knocked down, seen in 30 of the 66 tadpoles (45%) scored at this stage. The forelimb has not failed to hatch from its pocket, rather, it is completely absent (Figure 3A). Tadpole swimming is often hampered by the asymmetric nature of the head in *Recql4* unilateral CRISPants, which may lead to slower growth (<u>Video S1</u>). The endpoint of stage NF58 was chosen to avoid potential issues with mobility, as the tadpoles start to rely more on limbs and less on the tail as metamorphosis proceeds. However, around half of the CRISPR tadpoles were unable to reach stage NF58 several weeks after the last control (GFP) sibling was fixed, indicating that developmental delay is an issue even in half CRISPants. We therefore likely captured the weakest phenotypes, as they reached the goal stage of development earlier. Unexpectedly, three individuals developed ectopic forelimbs on the edited side (Figure 3B). In two cases, we saw mirror image duplicates. Figure 3B shows a right-sided CRISPant with two perfectly formed forelimbs joined at the shoulder and with mirror image polarity. In the third case, we saw a chaotic limb, the structure of which was revealed by skeletal staining (Figure 3B, left CRISPant). The limb contained at least three branches, each of which had one or more phalanges, but none of which was correctly patterned. Overall, limb defects were almost exclusively found on the CRISPR edited side of the tadpole, with only a single individual also lacking the contralateral forelimb (Figure 3C).

**Figure 3:**
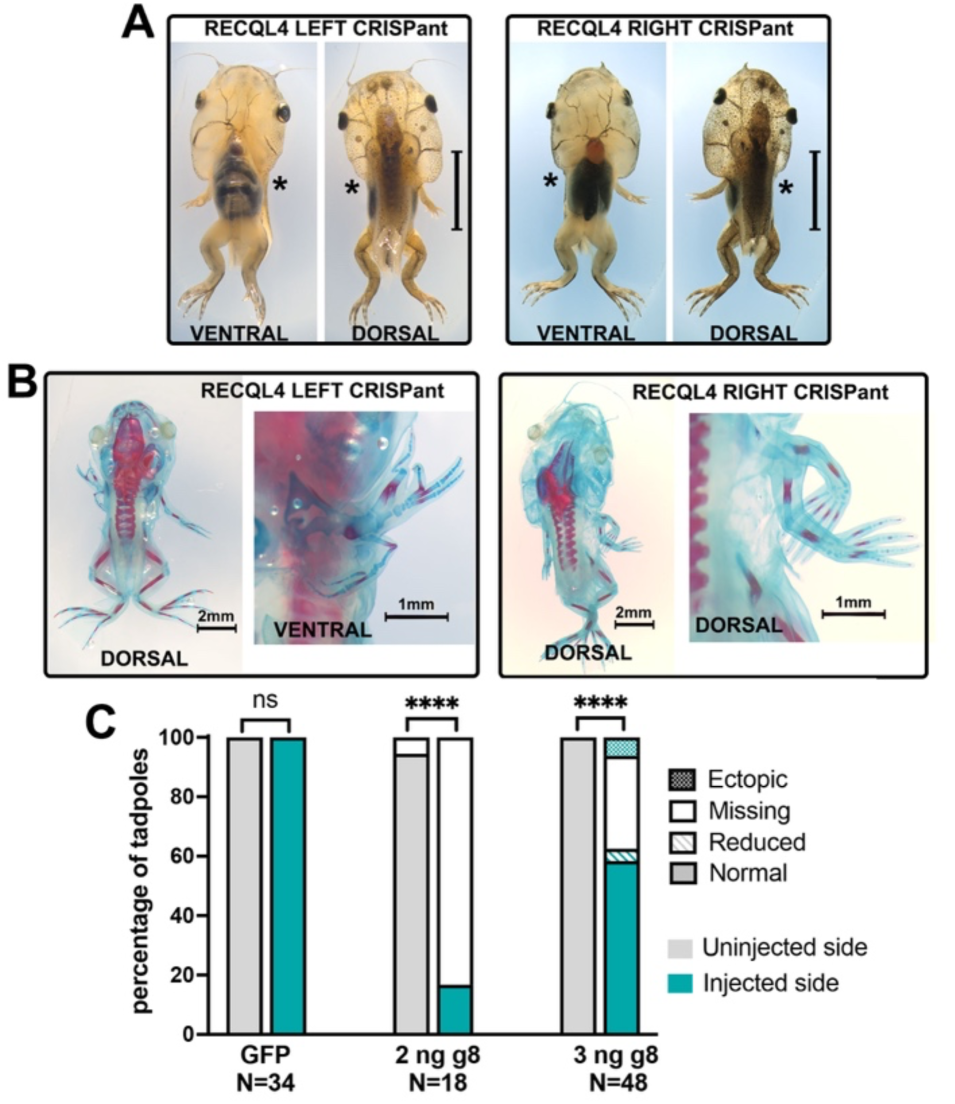
Forelimb development requires Recql4. **A)** Stage NF 58 tadpoles commonly lack a forelimb on the CRISPR edited side. Examples of left and right sided *Recql4* CRISPANT tadpoles, dorsal and ventral views. Missing forelimbs are indicated by an asterisk; scale bar is 5 mm. Tails have been removed to aid visualisation. **B)** Whole skeletal preparations of example stage NF 58 tadpoles where the edited side forelimb was not missing but instead was severely disrupted or duplicated, with ectopic structures present. Blue stain is cartilage (alcian blue) and red stain shows bone (alizarin red). The whole tadpole is shown in the left image and a zoomed view of the ectopic limb on the right, viewed from the side that best shows the limb elements. **C)** Stacked bar graph showing observed frequency of forelimb phenotypes on the uninjected (grey) side vs. injected (teal) side. Fishers exact test, ns= non-significant, **** = p<0.0001. Raw count data can be found in Table S7 in File S1 and images in Figure S2.

Hindlimbs were also significantly affected, although at a lower frequency, with 9/66 (13.6%) of skeletal preparations missing a hindlimb on the edited side and a further 9 with reduced limbs on the edited side (Figure 4, Table S7 in File S1) A single ectopic hindlimb was observed (Figure 4C) with mirror image duplication of the limb fused together at stylopod and zeugopod level by skin, and preaxial polydactyly of the autopod with 8 digits clearly visible. Skeletal staining of this individual revealed a single femur with duplicated tibia/fibula and two complete autopods, although both digit 1 copies had no ossification (Figure S4).

**Figure 4:**
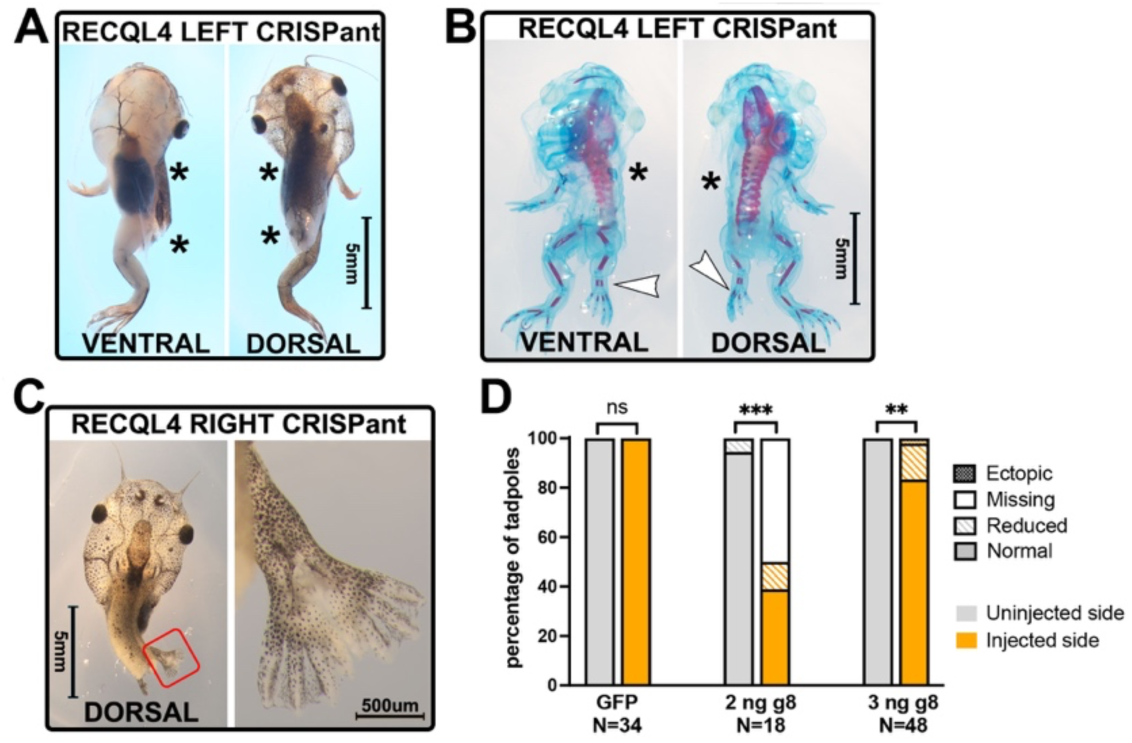
Hindlimbs are more resistant to loss of *Recql4* than forelimbs. **A)** Example of NF 58 stage tadpole where CRISPR knockdown of *recql4* on the left side has led to failure of the left hindlimb to develop as well as the forelimbs. Missing limbs are indicated by an asterisk. **B)** Skeletal preparation of NF58 tadpole showing example of a smaller hindlimb on the recql4 edited left side (white arrowhead). Bone and cartilage staining show that this limb is not missing any elements. This individual is also missing the left forelimb as indicated with an asterisk. **C)** A single example of an ectopic hindlimb found in the study is shown. This tadpole is stage NF55 and editing is on the right. Two limbs appear as a mirror image duplication (preaxial polydactyly) on the right side, and a single limb (partially obscured by the tail) on the left. Both forelimbs are present but cannot be seen due to the pre-hatching stage. **D)** Stacked bar graph showing observed frequency of hindlimb phenotypes on the injected vs. uninjected side of half-CRISPant tadpoles. Fisher’s exact tests, ns= non-significant, *** = p<0.001, ** p<0.01. Count data is in Table S7 in File S1.

### *Recql4* unilateral CRISPants have craniofacial defects on the edited side

The skeletal preparations used to examine limb development also provided an opportunity to observe craniofacial development and ossification. In total, we scored 52 unilateral CRISPant tadpoles for defects in ossification of the Meckel’s Cartilage (Figure 5). Ossified bone was absent on the CRISPR side in 42.3% of tadpoles and a further 36.5% had less ossified bone, or fragmentation of the ossified region, compared to the wild-type side (Figure 5A-C, Figure S3). This was clearly visible as reduced or absent alizarin red staining viewed from the ventral side. Other craniofacial structures were also affected, particularly the hyoid cartilage, which was smaller on the edited side in 71% of tadpoles, and completely missing in 1 individual (Figure 5A, B, D, Figure S3).

**Figure 5:**
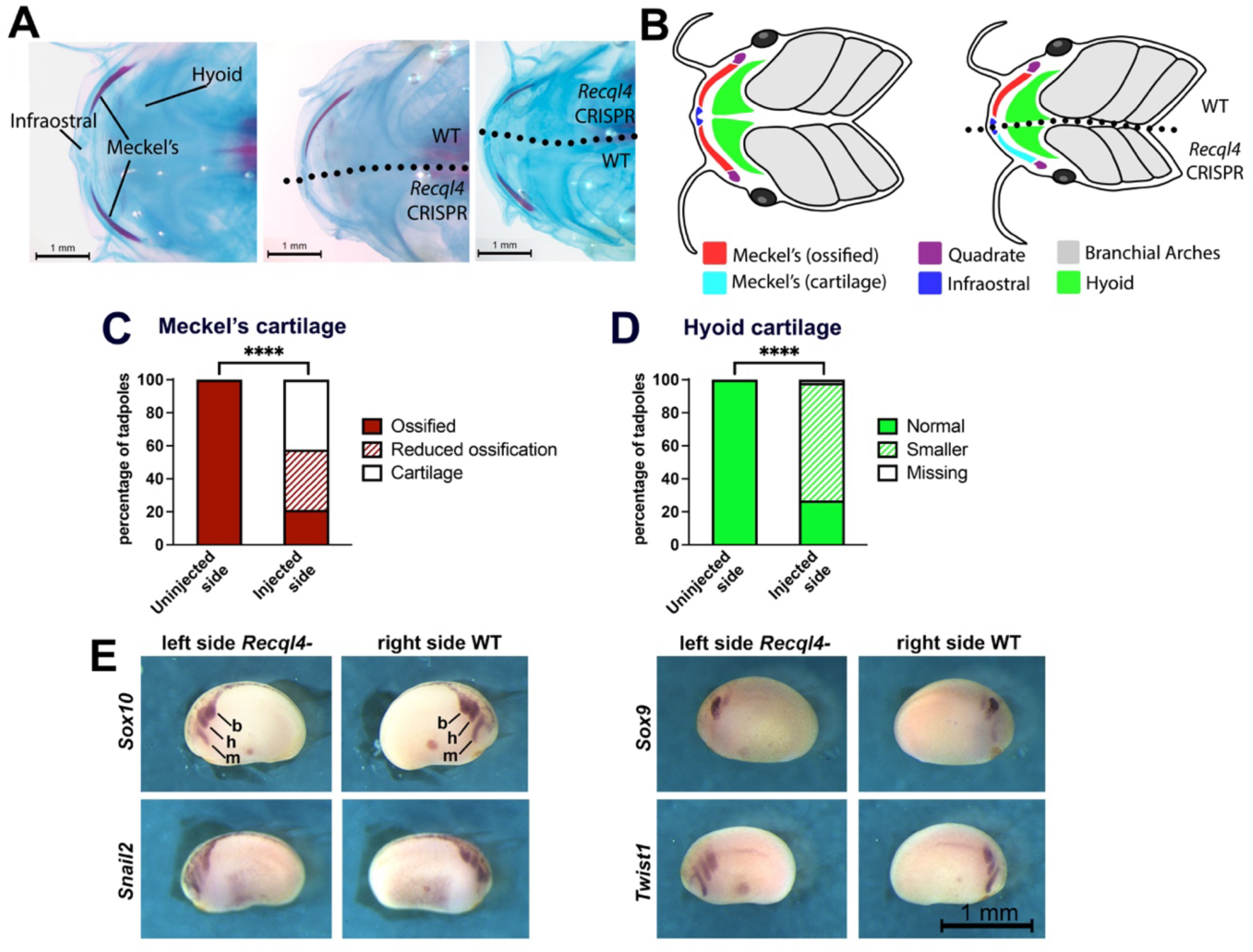
*Recql4* unilateral CRISPants have craniofacial defects on the edited side, but this does not arise from altered migration of cranial neural crest cells. A) Tadpole stage NF58 skeletal preparations, showing a ventral view of the jaw with cartilage stained blue and ossified bone stained red. Left image shows a control tadpole, centre shows a tadpole where *recql4* was knocked down on the left side (no ossification of Meckel’s cartilage on injected side), and right shows a tadpole with *recql4* knockdown on the right side (reduced ossification of Meckel’s on injected side. **B)** Diagram of control and left-sided CRISPant tadpole ventral craniofacial structures to show the positions of the cartilages. **C)** Stacked bar graph comparing the frequency of reduced or absent ossification of Meckel’s cartilage on the uninjected vs. injected side of unilateral *recql4* CRISPants. N=52, Fisher’s exact test p<0.0001 ****. **D)** Stacked bar graph comparing the frequency of smaller sized hyoid cartilage on the uninjected vs. injected unilateral CRISPants. N=52, Fisher’s exact test p<0.0001 ****. **E)** In situ hybridisation of early cranial neural crest markers in stage 19 unilateral *recql4* CRISPants (left side edited). The cranial neural crest cells migrate from the dorsal neural tube in three streams, the mandibular (m), hyoid (h) and branchial (b). The scale bar in the bottom right panel relates to all panels. Further examples can be found in Figure S3.

The cartilages of the head are built from the descendants of migrating cranial neural crest cells, which leave the dorsal neural tube in three streams during the neurula stages of development. Fate mapping experiments have shown that the most anterior of these, the mandibular stream, contributes to Meckel’s cartilage (lower jaw), quadrate cartilage (upper jaw) and parts of the skull, with the central stream giving rise to the hyoid cartilage (Sadaghiani & Thiébaud, 1987). Craniofacial defects like these are often the result of aberrant migration of neural crest, and *Xenopus* is a well-established model to determine this (reviewed in (Dubey & Saint-Jeannet, 2017). To see if the defects were caused by failure of these cells to migrate correctly when *recql4* has been edited, we examined the expression of four marker genes: *sox10, snail2, sox9* and *twist1* (Alfandari *et al*., 2001, Aoki *et al*., 2003, Lee & Saint-Jeannet, 2011) in unilateral *recql4* CRISPants (sgRNA8, Figure 5E). We saw no differences in the extent of migration or intensity of signal indicating that cranial neural crest migrates normally in the absence of Recql4. These results suggest that the reduced size of the hyoid cartilage results from hypoplastic growth.

### Forelimb buds fail to develop in the absence of Recql4, and vasculature of the head is hypoplastic with reduced branching

To determine whether the anterior limb buds developed and then regressed, or just never developed in the absence of Recql4, we examined tadpoles at four weeks of development using two different sgRNAs (Figure 6, Tables S8-S11 in File S1). Forelimb buds were absent on the ipsilateral side in 91.7% of tadpoles injected with sgRNA 8 (N=12), and 84.6% with sgRNA10 (N=26) (Figure 6A). The contralateral wild type forelimb was missing as well in a single sgRNA8 (8.3%) and a single sgRNA10 (3.8%).. Hindlimb defects were less commonly observed, but still missing on the ipsilateral side of 21.1% of *recql4* half-CRISPants (Figure 6A). Two individuals had had hindlimbs that were developmentally delayed on the ipsilateral side by two NF stagesJust two of 38 tadpoles had a missing hindlimb bud on the contralateral side (5%). These observations from an earlier timepoint in development support the conclusion that forelimb buds fail to develop in the absence of Recql4.

**Figure 6:**
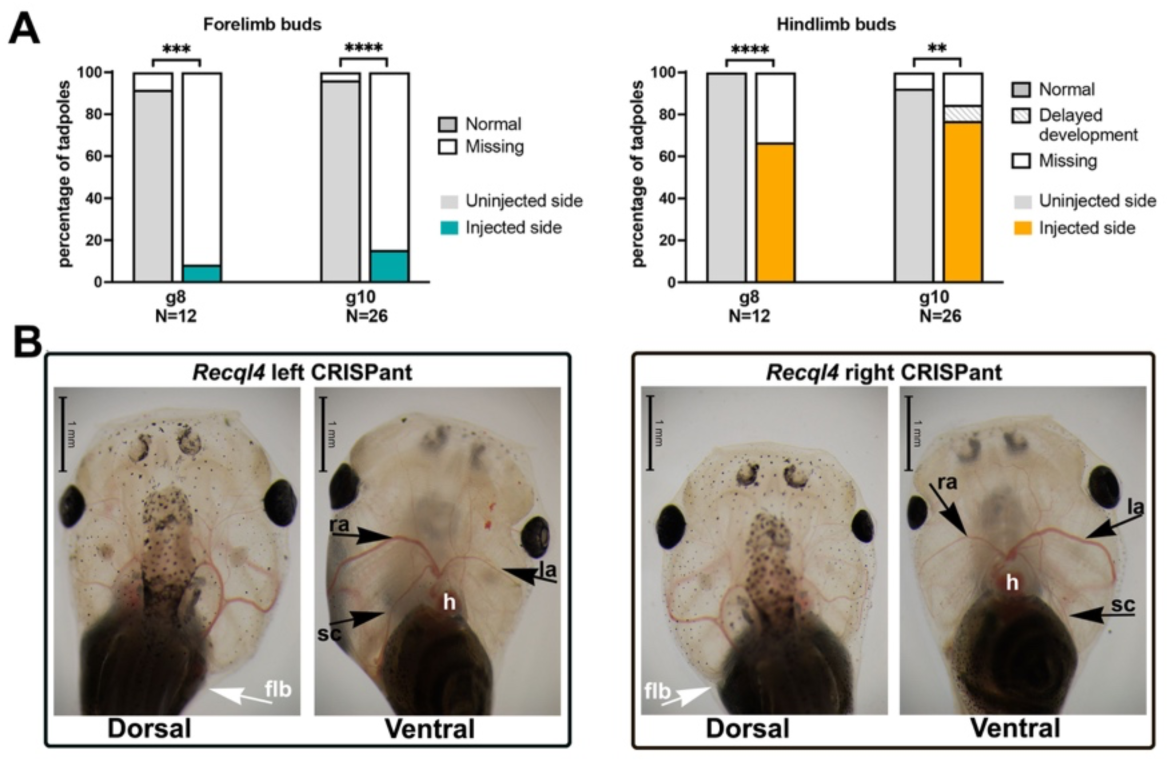
The forelimb bud fails to develop and vasculature of the head is reduced in the absense of Recql4. **A)** Stacked bar graphs showing observed frequency of forelimb absence or delayed development on uninjected vs. injected sides of half-CRISPant tadpoles at four weeks old (stages NF50-54). Fisher’s exact test, ** p<0.01, *** p<0.001, **** p<0.0001.**B)** Dorsal and ventral views of the head of typical unilateral CRISPant tadpoles at four weeks of development. The heart and vascular system can be clearly seen due to the natural transparency of the tadpole head.Labels: flb, forelimb bud; h, heart; ra, right aorta; la, left aorta; sc, subclavian artery. Raw data can be found in Tables S8-S11 in File S1.

While scoring the limbs, we noticed that the vasculature was also altered in the ipsilateral head region, with the CRISPR side consistently having a narrower aorta. Additionally, some arteries that branch from the aorta were absent on the CRISPR side, particularly the subclavian artery (Figure 6B, <u>Video S2, Video S3).</u>

## Discussion

Here we have used CRISPR/Cas9 editing to study the effects of depleting Recql4, using two sgRNAs in different positions to help further understand developmental consequences caused by frameshift variants in different parts of the *recql4* gene. Following injection of the two different sgRNAs, CRISPants showed concordant phenotypes, despite the difference in position of editing. In other animal models, different phenotypes have been observed depending on the variant position. For example, mouse models harbouring either p.Arg347* or p.G522fs variants showed different effects on the protein (weak expression of truncated protein versus no expression, respectively) and the p.Arg347* variant caused a more severe skeletal phenotypes (Castillo-Tandazo *et al*., 2021). A missense variant which inactivates the helicase, p.K525A, showed no phenotype in a mouse model, although the skeletal system was not studied in detail (Castillo-Tandazo *et al*., 2019). The majority of variants identified in humans have also not been characterised in a cellular system, so trying to establish genotype-phenotype patterns or design precise animal models remains difficult.

Translational effects of variants in the *recql4* gene appear to be complex in all models studied (Castillo-Tandazo *et al*., 2019, Castillo-Tandazo *et al*., 2021). While some variants that cause premature stop codons, through either nonsense or frameshift variants, result in little protein expression (likely through nonsense-mediated decay pathways regulating the aberrant mRNA), others result in the production of a truncated protein, albeit at a reduced level (Castillo-Tandazo *et al*., 2021). An unusual, evolutionarily conserved feature of the *recql4* genes is the significant number of very short introns. Exactly how these might impact mRNA quality surveillance and splicing has not been determined, but these likely play a role in the complexity of Recql4 translational products and levels when variants are present.

A strong advantage to our approach was to harness the unique left/right separation of the body plan which is present in *X. laevis* from the very first cell division. This enables targeting of CRISPR editing to one side of the body, with the co-injection of mRNA for nuclear GFP allows accurate sorting of embryos from neurula stages (24 hours from injection) into left and right CRISPants. Amplicon Sanger sequencing showed that both guides were efficient editors, resulting in around 50% of the genome overall, indicting that levels of editing on the targeted side are close to 100%. As demonstrated, this enabled an inbuilt control and thereby a more precise study of phenotypes caused by gene editing. The injected side of the CRISPant tadpoles had smaller heads and eyes. Growth defects, including short stature and/or microcephaly, are well established in human disorders associated with *RECQL4* (Wang & Plon, 1999, Van Maldergem *et al*., 2007). Eyes are a highly proliferative tissue, and in other animal models where microcephaly has occurred, smaller eyes can also be present (Hempel *et al*., 2016, Johnson *et al*., 2024, Satou-Kobayashi *et al*., 2024). Microphthalmia has only been noted in one individual with *RECQL4* variants (Yadav *et al*., 2019) and so the observation that CRISPants have smaller eyes, is more likely a secondary consequence of overall reduced growth, rather than a distinct feature.

The skeletal defects caused by the editing aligned with the human cases of *RECQL4* and mouse skeletal models (Kitao *et al*., 1999, Siitonen *et al*., 2003, Piard *et al*., 2015). Forelimbs were much more likely to be absent rather than hypoplastic, suggesting a failure early in limb development. The effect on hindlimbs was not as commonly observed, but similar in that the limb was more likely to be absent rather than reduced. This could either reflect the editing approach used, or insight into the underlying biology. While forelimbs develop on opposite sides of the trunk, hindlimb buds develop very close together at the midline. Cell mixing across the midline due to this proximity could therefore contribute some wild type cells to the edited limb mesenchyme, giving a threshold amount of wildtype Recql4 on the edited sides. The forelimb buds are much further apart, so cell mixing would be less likely. However, it is noteworthy that in humans harbouring *RECQL4* variants, arms are also more often affected than legs and feet. Radial ray defects are a specific grouping of skeletal anomalies affecting the radius, radial carpal bones and thumb, arising from failures in limb bud mesoderm growth and patterning, but there is not an equivalent grouping affecting legs and feet. Forelimb absence in our CRISPants could be considered an extreme failure of limb bud mesoderm growth and patterning.

In a few cases we did observe ectopic hind or forelimbs developing, resembling complete mirror image polydactyly caused by mis-expression of *shh* (Riddle *et al*., 1993) or grafting of the zone of polarising activity ZPA (Honig & Summerbell, 1985) in chickens. A duplicated thumb (preaxial polydactyly) has been reported in a single case with *RECQL4* variants (Piard *et al*., 2015). While this unexpected phenotype has been observed in both *X. laevis* and humans, it is not clear how Recql4 knockdown could lead to these diametrically opposed phenotypes of hypomorphic/missing limbs and anterior-posterior limb duplications. Seemingly opposed phenotypes of limb hypoplasia and limb duplication have also been observed to result from overexpression of the BMP inhibitor Noggin1 at different developmental stages in inducible transgenic lines (Jones *et al*., 2013). These different observations further support that Recql4 must have specific role(s) in skeletal development and potentially form a starting point to understand the link between Recql4 function and developmental processes in the limb bud.

We also observed some features unique to our CRISPant models. While some facial dysmorphism is occasionally seen in *RECQL4*-associated individuals (Piard *et al*., 2015), including micrognathia, we observed stronger phenotypes in our models. The edited side of the head was smaller overall and often showed either reduced, fragmented or a total lack of ossification. The ossification of Meckel’s cartilage and the size of the hyoid cartilage were particularly sensitive to loss of Recql4 – although for the hyoid cartilage it was more likely reduced in the size than absent, which is different to the forelimb and hindlimb consequences we observed. Using in situ hybridization, we confirmed that this is not due to generalised neural crest migration defects, suggesting a more specific effect on these structures caused by reduced or absent Recql4. Meckel’s cartilage is a conserved developmental feature of all jawed vertebrates, which acts as a template for later forming jaw bones, but it has evolved a distinct role in mammals whereby the proximal part ossifies and forms the malleus and incus, two of the three middle ear bones (Svandova *et al*., 2020). There are no reports of disruption to these bones in the human *recql4* syndromes that we are aware of. It is also not clear whether the lack of ossification in stage 58 tadpoles is a delay in the process or complete failure, since tadpoles were not raised past this stage. Despite the differences in anatomy, the observations of delayed or failed ossification in both the tadpole Meckel’s cartilage (this study) and the human patella (Wang & Plon, 1999) suggest a role for Recql4 in endochondral ossification. Failure to ossify the patella also occurs in Meier-Gorlin syndrome, a multigenic disorder that results in growth retardation due to impaired replication licencing (Klingseisen & Jackson, 2011, de Munnik *et al*., 2015).

Intriguingly, when checking the younger stage tadpoles to see if limb buds ever formed, we observed absence of a major nearby blood vessel, likely the subclavian artery. No vasculature abnormalities have been noted in human patients, although links between angiogenesis (the formation of new blood vessels) and limb development are highlighted by the dramatic teratogenic nature of the drug thalidomide (reviewed in (Vargesson & Hootnick, 2017)). A failure to form both the forelimb bud and the subclavian artery that supplies the forelimb may suggest a developmental co-dependence. The timing of the formation of the *Xenopus* subclavian artery has not been described, but in humans and mice it seems to only become established *after* the initiation of limb bud development, (Rodríguez-Niedenführ *et al*., 2001, Walls *et al*., 2008). Alternatively, reduced aortic branching could result from interference with the VEGF signalling that drives this process, or simply reflect the poor growth on the side of the tadpole that has Recql4 knockdown. The aortic arch appeared consistently narrower on the CRISPR side, suggesting the latter.

In conclusion, our model establishes single-sided genome editing as a useful approach for studying developmental effects of reduced Recql4. This method could be employed to other monogenic disorders that associated with extreme growth (Bicknell *et al*., 2025). Many phenotypes observed in edited tadpoles align with both the human and mouse models, but we have also observed unique abnormalities that could either point to *X. laevis* specific requirements for Recql4, or broader effects caused by a developmental role for Recql4 that has not yet been fully determined. The cellular roles for RECQL4 continue to be explored (Ashraf *et al*., 2025); such further insight may aid in deciphering these unique phenotypes observed.

## Acknowledgements

We gratefully acknowledge funding from the Neurological Foundation of New Zealand (PRG1732 to LSB, CB). We would like to thank Nikita Woodhead and her team for her excellent care of the *Xenopus* colony.

## Author contributions

Experimental design, implementation and analysis CB, MRB. Animal ethics CB, Manuscript writing and revision CB, MRB, LSB.

## Data availability statement

All data are available in the main text or supplemental data file S1, and videos are available via FigShare.

## Supplementary data

File S1 contains raw data and statistical analysis tables S1-S11

Supplementary figures S1-S4 contain raw data and images.

DOI for supplementary videos 1-3, hosted at Figshare are as follows:

Video S1: DOI 10.6084/m9.figshare.28640594

Video S2: DOI 10.6084/m9.figshare.28639817

Video S3: DOI 10 10.6084/m9.figshare.28640423

## Notes

https://doi.org/10.6084/m9.figshare.29740736

### Competing Interest Statement

The authors have declared no competing interest.

### Summary of Updates

Revised version following peer review. Some data has been condensed (left and right sided CRISPants) affecting almost all figures from the original, but not changing the findings. Stats have been added to 3 figures. Additional data around the sgRNA 10 editing has been included.

## References

Abu-Libdeh B, Jhujh SS, Dhar S, et al. (2022) RECON syndrome is a genome instability disorder caused by mutations in the DNA helicase RECQL1. J Clin Invest 132: e147301.

Alfandari D, Cousin H, Gaultier A, Smith K, White JM, Darribère T & DeSimone DW (2001) Xenopus ADAM 13 is a metalloprotease required for cranial neural crest-cell migration. Curr Biol 11: 918–930.

Aoki Y, Saint-Germain N, Gyda M, Magner-Fink E, Lee YH, Credidio C & Saint-Jeannet JP (2003) Sox10 regulates the development of neural crest-derived melanocytes in Xenopus. Dev Biol 259: 19–33.

Arora H, Chacon AH, Choudhary S, McLeod MP, Meshkov L, Nouri K & Izakovic J (2014) Bloom syndrome. Int J Dermatol 53: 798–802.

Ashraf R, Polasek-Sedlackova H, Marini V, Prochazkova J, Hasanova Z, Zacpalova M, Boudova M & Krejci L (2025) RECQ4-MUS81 interaction contributes to telomere maintenance with implications to Rothmund-Thomson syndrome. Nature Communications 16: 1302.

Beck CW & Slack JM (1998) Analysis of the developing Xenopus tail bud reveals separate phases of gene expression during determination and outgrowth. Mech Dev 72: 41–52.

Bellelli R & Boulton SJ (2021) Spotlight on the Replisome: Aetiology of DNA Replication-Associated Genetic Diseases. Trends in Genetics 37: 317–336.

Bicknell LS, Hirschhorn JN & Savarirayan R (2025) The genetic basis of human height. Nat Rev Genet.

Bohr VA (2008) Rising from the RecQ-age: the role of human RecQ helicases in genome maintenance. Trends Biochem Sci 33: 609–620.

Brinkman EK, Chen T, Amendola M & van Steensel B (2014) Easy quantitative assessment of genome editing by sequence trace decomposition. Nucleic Acids Res 42: e168.

Capp C, Wu J & Hsieh TS (2009) Drosophila RecQ4 has a 3’-5’ DNA helicase activity that is essential for viability. J Biol Chem 284: 30845–30852.

Castillo-Tandazo W, Frazier AE, Sims NA, Smeets MF & Walkley CR (2021) Rothmund-Thomson Syndrome-Like RECQL4 Truncating Mutations Cause a Haploinsufficient Low-Bone-Mass Phenotype in Mice. Mol Cell Biol 41: e0059020.

Castillo-Tandazo W, Smeets MF, Murphy V, Liu R, Hodson C, Heierhorst J, Deans AJ & Walkley CR (2019) ATP-dependent helicase activity is dispensable for the physiological functions of Recql4. PLOS Genetics 15: e1008266.

Chapman PA, Gilbert CB, Devine TJ, Hudson DT, Ward J, Morgan XC & Beck CW (2022) Manipulating the microbiome alters regenerative outcomes in Xenopus laevis tadpoles via lipopolysaccharide signalling. Wound Repair Regen 30: 636–651.

Croteau DL, Popuri V, Opresko PL & Bohr VA (2014) Human RecQ helicases in DNA repair, recombination, and replication. Annu Rev Biochem 83: 519–552.

de Munnik SA, Hoefsloot EH, Roukema J, Schoots J, Knoers NV, Brunner HG, Jackson AP & Bongers EM (2015) Meier-Gorlin syndrome. Orphanet J Rare Dis 10: 114.

Dubey A & Saint-Jeannet JP (2017) Modeling human craniofacial disorders in Xenopus. Curr Pathobiol Rep 5: 79–92.

Fisher M, James-Zorn C, Ponferrada V, et al. (2023) Xenbase: key features and resources of the Xenopus model organism knowledgebase. Genetics 224.

Hempel A, Pagnamenta AT, Blyth M, et al. (2016) Deletions and de novo mutations of SOX11 are associated with a neurodevelopmental disorder with features of Coffin-Siris syndrome. J Med Genet 53: 152–162.

Hickson ID (2003) RecQ helicases: caretakers of the genome. Nature Reviews Cancer 3: 169–178.

Hoki Y, Araki R, Fujimori A, et al. (2003) Growth retardation and skin abnormalities of the Recql4 -deficient mouse. Human Molecular Genetics 12: 2293–2299.

Honig LS & Summerbell D (1985) Maps of strength of positional signalling activity in the developing chick wing bud. J Embryol Exp Morphol 87: 163–174.

Ichikawa K, Noda T & Furuichi Y (2002) [Preparation of the gene targeted knockout mice for human premature aging diseases, Werner syndrome, and Rothmund-Thomson syndrome caused by the mutation of DNA helicases]. Nihon Yakurigaku Zasshi 119: 219–226.

Jiang C, Zhang H, Zhao C, Wang L, Hu X & Pan Z (2023) De novo myelodysplastic syndrome in a Rothmund-Thomson Syndrome patient with novel pathogenic RECQL4 variants. Blood Sci 5: 125–130.

Johnson HK, Wahl SE, Sesay F, Litovchick L & Dickinson AJ (2024) Dyrk1a is required for craniofacial development in Xenopus laevis. Dev Biol 511: 63–75.

Jones TE, Day RC & Beck CW (2013) Attenuation of bone morphogenetic protein signaling during amphibian limb development results in the generation of stage-specific defects. J Anat 223: 474–488.

Kaiser S, Sauer F & Kisker C (2017) The structural and functional characterization of human RecQ4 reveals insights into its helicase mechanism. Nature Communications 8: 15907.

Kitao S, Shimamoto A, Goto M, Miller RW, Smithson WA, Lindor NM & Furuichi Y (1999) Mutations in RECQL4 cause a subset of cases of Rothmund-Thomson syndrome. Nat Genet 22: 82–84.

Klingseisen A & Jackson AP (2011) Mechanisms and pathways of growth failure in primordial dwarfism. Genes Dev 25: 2011–2024.

Labun K, Montague TG, Krause M, Torres Cleuren YN, Tjeldnes H & Valen E (2019) CHOPCHOP v3: expanding the CRISPR web toolbox beyond genome editing. Nucleic Acids Research 47: W171–W174.

Larizza L, Roversi G & Volpi L (2010) Rothmund-Thomson syndrome. Orphanet J Rare Dis 5: 2.

Larsen NB & Hickson ID (2013) RecQ Helicases: Conserved Guardians of Genomic Integrity. Adv Exp Med Biol 767: 161–184.

Lee YH & Saint-Jeannet JP (2011) Sox9 function in craniofacial development and disease. Genesis 49: 200–208.

Lu H, Shamanna RA, de Freitas JK, et al. (2017) Cell cycle-dependent phosphorylation regulates RECQL4 pathway choice and ubiquitination in DNA double-strand break repair. Nature Communications 8: 2039.

Lu L, Harutyunyan K, Jin W, et al. (2015) RECQL4 Regulates p53 Function In Vivo During Skeletogenesis. J Bone Miner Res 30: 1077-1089.

Luong TT & Bernstein KA (2021) Role and Regulation of the RECQL4 Family during Genomic Integrity Maintenance. Genes (Basel) 12.

Mann MB, Hodges CA, Barnes E, Vogel H, Hassold TJ & Luo G (2005) Defective sister-chromatid cohesion, aneuploidy and cancer predisposition in a mouse model of type II Rothmund-Thomson syndrome. Hum Mol Genet 14: 813–825.

Martins DJ, Di Lazzaro Filho R, Bertola DR & Hoch NC (2023) Rothmund-Thomson syndrome, a disorder far from solved. Frontiers in Aging 4.

Mojumdar A, De March M, Marino F & Onesti S (2017) The Human RecQ4 Helicase Contains a Functional RecQ C-terminal Region (RQC) That Is Essential for Activity. J Biol Chem 292: 4176–4184.

Moreno-Mateos MA, Vejnar CE, Beaudoin JD, Fernandez JP, Mis EK, Khokha MK & Giraldez AJ (2015) CRISPRscan: designing highly efficient sgRNAs for CRISPR-Cas9 targeting in vivo. Nat Methods 12: 982–988.

Nenni MJ, Fisher ME, James-Zorn C, et al. (2019) Xenbase: Facilitating the Use of Xenopus to Model Human Disease. Front Physiol 10: 154.

Newman SM, Jr. & Dumont JN (1983) Thiosemicarbazide-induced osteolathyrism in metamorphosing Xenopus laevis. J Exp Zool 225: 411–421.

Nieuwkoop PD & Faber J (1994) Normal Table of Xenopus laevis (Daudin): a systematical and chronological survey of the development from the fertilized egg till the end of metamorphosis. Garland Publishing Inc, New York & London.

Oshima J, Sidorova JM & Monnat RJ, Jr. (2017) Werner syndrome: Clinical features, pathogenesis and potential therapeutic interventions. Ageing Res Rev 33: 105–114.

Piard J, Aral B, Vabres P, et al. (2015) Search for ReCQL4 mutations in 39 patients genotyped for suspected Rothmund-Thomson/Baller-Gerold syndromes. Clin Genet 87: 244–251.

Riddle RD, Johnson RL, Laufer E & Tabin C (1993) Sonic hedgehog mediates the polarizing activity of the ZPA. Cell 75: 1401–1416.

Rodríguez-Niedenführ M, Burton GJ, Deu J & Sañudo JR (2001) Development of the arterial pattern in the upper limb of staged human embryos: normal development and anatomic variations. J Anat 199: 407–417.

Sadaghiani B & Thiébaud CH (1987) Neural crest development in the Xenopus laevis embryo, studied by interspecific transplantation and scanning electron microscopy. Developmental Biology 124: 91–110.

Sangrithi MN, Bernal JA, Madine M, Philpott A, Lee J, Dunphy WG & Venkitaraman AR (2005) Initiation of DNA replication requires the RECQL4 protein mutated in Rothmund-Thomson syndrome. Cell 121: 887–898.

Satou-Kobayashi Y, Takahashi S, Haramoto Y, Asashima M & Taira M (2024) Zbtb11 interacts with Otx2 and patterns the anterior neuroectoderm in Xenopus. PLOS ONE 19: e0293852.

Schindelin J, Arganda-Carreras I, Frise E, et al. (2012) Fiji: an open-source platform for biological-image analysis. Nature Methods 9: 676-682.

Shen MW, Arbab M, Hsu JY, Worstell D, Culbertson SJ, Krabbe O, Cassa CA, Liu DR, Gifford DK & Sherwood RI (2018) Predictable and precise template-free CRISPR editing of pathogenic variants. Nature 563: 646–651.

Siitonen HA, Kopra O, Kääriäinen H, Haravuori H, Winter RM, Säämänen AM, Peltonen L & Kestilä M (2003) Molecular defect of RAPADILINO syndrome expands the phenotype spectrum of RECQL diseases. Hum Mol Genet 12: 2837–2844.

Siitonen HA, Sotkasiira J, Biervliet M, et al. (2009) The mutation spectrum in RECQL4 diseases. European Journal of Human Genetics 17: 151–158.

Singh DK, Karmakar P, Aamann M, Schurman SH, May A, Croteau DL, Burks L, Plon SE & Bohr VA (2010) The involvement of human RECQL4 in DNA double-strand break repair. Aging Cell 9: 358–371.

Smeets MF, DeLuca E, Wall M, Quach JM, Chalk AM, Deans AJ, Heierhorst J, Purton LE, Izon DJ & Walkley CR (2014) The Rothmund-Thomson syndrome helicase RECQL4 is essential for hematopoiesis. The Journal of Clinical Investigation 124: 3551–3565.

Svandova E, Anthwal N, Tucker AS & Matalova E (2020) Diverse Fate of an Enigmatic Structure: 200 Years of Meckel’s Cartilage. Front Cell Dev Biol 8: 821.

Van Maldergem L, Piard J, Larizza L & Wang LL (2007) Baller Gerold Syndrome. GeneReviews,(P. AM, J. F & M. MG, eds.), p.^pp. University of Washington, Seattle, WA, USA.

Van Maldergem L, Piard J, Larizza L & Wang LL (2016) 1141RECQL4-Related Recessive Conditions. Epstein’s Inborn Errors of Development: The Molecular Basis of Clinical Disorders of Morphogenesis, (Erickson RP & Wynshaw-Boris AJ, eds.), p.^pp. 0. Oxford University Press.

Van Maldergem L, Siitonen HA, Jalkh N, et al. (2006) Revisiting the craniosynostosis-radial ray hypoplasia association: Baller-Gerold syndrome caused by mutations in the RECQL4 gene. J Med Genet 43: 148–152.

Vargesson N & Hootnick DR (2017) Arterial dysgenesis and limb defects: Clinical and experimental examples. Reprod Toxicol 70: 21–29.

Walls JR, Coultas L, Rossant J & Henkelman RM (2008) Three-dimensional analysis of vascular development in the mouse embryo. PLoS One 3: e2853.

Wang LL & Plon SE (1999) Rothmund-Thomson Syndrome. Gene Reviews,(P. AM, J. F & M. MG, eds.), p.^pp. University of Washington, Seattle, WA, USA.

Wang LL, Gannavarapu A, Kozinetz CA, et al. (2003) Association between osteosarcoma and deleterious mutations in the RECQL4 gene in Rothmund-Thomson syndrome. J Natl Cancer Inst 95: 669–674.

Willsey HR, Seaby EG, Godwin A, Ennis S, Guille M & Grainger RM (2024) Modelling human genetic disorders in Xenopus tropicalis. Dis Model Mech 17.

Willsey HR, Exner CRT, Xu Y, et al. (2021) Parallel in vivo analysis of large-effect autism genes implicates cortical neurogenesis and estrogen in risk and resilience. Neuron 109: 1409.

Yadav S, Thakur S, Kohlhase J, Bhari N, Kabra M & Gupta N (2019) Report of Two Novel Mutations in Indian Patients with Rothmund-Thomson Syndrome. J Pediatr Genet 8: 163–167.

