## Supplementary material for "Unilateral loss of *recql4* function in *Xenopus laevis* tadpoles leads to ipsilateral ablation of the forelimb, hypoplastic Meckel’s cartilage and vascular defects": File S1

**Supplementary Information**

**Table S1:Left CRISPant head and eye measurements at 21 days old.**

| **Left edited**  **tadpole image** | **Area (mm^2^)** | | | | **Ratio injected/non-injected side** | |
| --- | --- | --- | --- | --- | --- | --- |
|  | **Left head** | **Right head** | **left eye** | **right eye** | **head** | **Eye** |
| control_left1 | 6.389 | 5.184 | 0.286 | 0.256 | 1.232 | 1.117 |
| control_left2 | 9.962 | 11.048 | 0.423 | 0.378 | 0.902 | 1.119 |
| control_left3 | 9.705 | 8.506 | 0.319 | 0.316 | 1.141 | 1.009 |
| control_left4 | 8.289 | 8.743 | 0.372 | 0.354 | 0.948 | 1.051 |
| control_left5 | 8.936 | 9.595 | 0.405 | 0.398 | 0.931 | 1.018 |
| control_left6 | 9.238 | 7.831 | 0.326 | 0.361 | 1.180 | 0.903 |
| control_left7 | 7.900 | 8.553 | 0.340 | 0.339 | 0.924 | 1.003 |
| control_left8 | 7.749 | 8.081 | 0.349 | 0.355 | 0.959 | 0.983 |
| control_left9 | 9.563 | 9.696 | 0.380 | 0.384 | 0.986 | 0.990 |
| control_left10 | 6.984 | 7.812 | 0.337 | 0.341 | 0.894 | 0.988 |
| control_left11 | 7.518 | 7.535 | 0.330 | 0.309 | 0.998 | 1.068 |
| control_left12 | 5.605 | 5.620 | 0.277 | 0.262 | 0.997 | 1.057 |
| control_left13 | 10.909 | 9.622 | 0.352 | 0.360 | 1.134 | 0.978 |
| control_left14 | 6.346 | 6.469 | 0.301 | 0.300 | 0.981 | 1.003 |
| control_left15 | 8.996 | 9.239 | 0.395 | 0.382 | 0.974 | 1.034 |
| Recql4_g8_2ng_left1 | 3.871 | 7.053 | 0.206 | 0.296 | 0.549 | 0.696 |
| Recql4_g8_2ng_left2 | 5.291 | 6.847 | 0.333 | 0.349 | 0.773 | 0.954 |
| Recql4_g8_2ng_left3 | 4.195 | 6.321 | 0.310 | 0.310 | 0.664 | 1.000 |
| Recql4_g8_2ng_left4 | 3.141 | 8.451 | 0.268 | 0.387 | 0.372 | 0.693 |
| Recql4_g8_2ng_left5 | 2.975 | 4.196 | 0.134 | 0.237 | 0.709 | 0.565 |
| Recql4_g8_2ng_left6 | 4.934 | 6.473 | 0.257 | 0.280 | 0.762 | 0.918 |
| Recql4_g8_2ng_left7 | 6.224 | 8.666 | 0.400 | 0.458 | 0.718 | 0.873 |
| Recql4_g8_2ng_left8 | 3.906 | 5.947 | 0.247 | 0.257 | 0.657 | 0.961 |
| Recql4_g8_2ng_left9 | 3.901 | 5.948 | 0.209 | 0.323 | 0.656 | 0.647 |
| Recql4_g8_2ng_left10 | 1.891 | 2.440 | 0.053 | 0.146 | 0.775 | 0.363 |
| Recql4_g8_2ng_left11 | 3.122 | 5.268 | 0.115 | 0.235 | 0.593 | 0.489 |
| Recql4_g8_3ng_left1 | 4.149 | 4.630 | 0.238 | 0.240 | 0.896 | 0.992 |
| Recql4_g8_3ng_left2 | 5.635 | 5.142 | 0.248 | 0.202 | 1.096 | 1.228 |
| Recql4_g8_3ng_left3 | 3.828 | 5.554 | 0.185 | 0.324 | 0.689 | 0.571 |
| Recql4_g8_3ng_left4 | 5.123 | 6.187 | 0.257 | 0.299 | 0.828 | 0.860 |
| Recql4_g8_3ng_left5 | 4.187 | 6.454 | 0.205 | 0.299 | 0.649 | 0.686 |
| Recql4_g8_3ng_left6 | 3.442 | 5.459 | 0.155 | 0.268 | 0.631 | 0.578 |
| Recql4_g8_3ng_left7 | 3.634 | 5.293 | 0.21 | 0.243 | 0.687 | 0.864 |
| Recql4_g8_3ng_left8 | 4.035 | 6.143 | 0.198 | 0.273 | 0.657 | 0.725 |
| Recql4_g8_3ng_left9 | 4.868 | 5.825 | 0.249 | 0.251 | 0.836 | 0.992 |
| Recql4_g8_3ng_left10 | 5.697 | 7.565 | 0.264 | 0.279 | 0.753 | 0.946 |
| Recql4_g8_3ng_left11 | 3.823 | 5.147 | 0.216 | 0.239 | 0.743 | 0.904 |
| Recql4_g8_3ng_left12 | 3.455 | 3.884 | 0.194 | 0.201 | 0.890 | 0.965 |
| Recql4_g10_left1 | 4.444 | 5.855 | 0.202 | 0.248 | 0.759 | 0.815 |
| Recql4_g10_left2 | 3.716 | 6.132 | 0.168 | 0.266 | 0.606 | 0.632 |
| Recql4_g10_left3 | 4.028 | 4.412 | 0.157 | 0.228 | 0.913 | 0.689 |
| Recql4_g10_left4 | 3.646 | 5.696 | 0.162 | 0.239 | 0.640 | 0.678 |
| Recql4_g10_left5 | 2.889 | 4.195 | 0.072 | 0.196 | 0.689 | 0.367 |
| Recql4_g10_left6 | 2.789 | 3.308 | 0.057 | 0.167 | 0.843 | 0.341 |
| Recql4_g10_left7 | 2.176 | 3.446 | 0.076 | 0.157 | 0.631 | 0.484 |
| Recql4_g10_left8 | 1.514 | 1.783 | 0.047 | 0.122 | 0.849 | 0.385 |
| Recql4_g10_left9 | 3.042 | 4.635 | 0.141 | 0.228 | 0.656 | 0.618 |
| Recql4_g10_left10 | 2.174 | 2.547 | 0.056 | 0.131 | 0.854 | 0.427 |
| Recql4_g10_left11 | 2.043 | 2.972 | 0.078 | 0.164 | 0.687 | 0.476 |
| Recql4_g10_left12 | 1.859 | 2.121 | 0.098 | 0.100 | 0.876 | 0.980 |
| Recql4_g10_left13 | 1.685 | 1.990 | 0.066 | 0.106 | 0.847 | 0.623 |
| Recql4_g10_left14 | 1.533 | 1.553 | 0.061 | 0.075 | 0.987 | 0.813 |
| Recql4_g10_left15 | 1.488 | 2.057 | 0.048 | 0.135 | 0.723 | 0.356 |
| Recql4_g10_left16 | 1.310 | 1.551 | 0.075 | 0.094 | 0.845 | 0.798 |

**Table S2: Right CRISPant head and eye measurements at 21 days old.**

| **Right edited**  **tadpole image** | **Area (mm^2^)** | | | | **Ratio injected/non-injected side** | |
| --- | --- | --- | --- | --- | --- | --- |
|  | **Left head** | **Right head** | **left eye** | **right eye** | **head** | **eye** |
| control_right1 | 9.292 | 10.279 | 0.403 | 0.465 | 1.106 | 1.154 |
| control_right2 | 8.026 | 9.692 | 0.382 | 0.384 | 1.208 | 1.005 |
| control_right3 | 6.966 | 7.836 | 0.3 | 0.331 | 1.125 | 1.103 |
| control_right4 | 12.571 | 14.131 | 0.462 | 0.485 | 1.124 | 1.050 |
| control_right5 | 9.34 | 11.374 | 0.319 | 0.368 | 1.218 | 1.154 |
| control_right6 | 12.207 | 12.235 | 0.38 | 0.441 | 1.002 | 1.161 |
| control_right7 | 10.393 | 9.378 | 0.345 | 0.377 | 0.902 | 1.093 |
| control_right8 | 13.348 | 13.33 | 0.452 | 0.448 | 0.999 | 0.991 |
| control_right9 | 7.576 | 8.049 | 0.347 | 0.379 | 1.062 | 1.092 |
| control_right10 | 7.847 | 8.928 | 0.353 | 0.366 | 1.138 | 1.037 |
| control_right11 | 8.284 | 8.925 | 0.336 | 0.329 | 1.077 | 0.979 |
| control_right12 | 6.953 | 6.056 | 0.311 | 0.329 | 0.871 | 1.058 |
| control_right13 | 2.372 | 2.583 | 0.126 | 0.145 | 1.089 | 1.151 |
| Recql4_g8_2ng_right1 | 4.467 | 3.801 | 0.272 | 0.287 | 0.851 | 1.055 |
| Recql4_g8_2ng_right2 | 3.626 | 3.12 | 0.271 | 0.278 | 0.860 | 1.026 |
| Recql4_g8_2ng_right3 | 3.325 | 2.193 | 0.169 | 0.045 | 0.660 | 0.266 |
| Recql4_g8_2ng_right4 | 4.175 | 2.68 | 0.215 | 0.102 | 0.642 | 0.474 |
| Recql4_g8_2ng_right5 | 3.31 | 2.354 | 0.17 | 0.07 | 0.711 | 0.412 |
| Recql4_g8_2ng_right6 | 5.842 | 5.33 | 0.272 | 0.256 | 0.912 | 0.941 |
| Recql4_g8_2ng_right7 | 4.088 | 3.133 | 0.259 | 0.129 | 0.766 | 0.498 |
| Recql4_g8_2ng_right8 | 6.068 | 4.51 | 0.277 | 0.292 | 0.743 | 1.054 |
| Recql4_g8_2ng_right9 | 4.417 | 2.974 | 0.229 | 0.229 | 0.673 | 1.000 |
| Recql4_g8_3ng_right1 | 7.740 | 5.398 | 0.36 | 0.302 | 0.697 | 0.839 |
| Recql4_g8_3ng_right2 | 9.628 | 7.275 | 0.333 | 0.306 | 0.756 | 0.919 |
| Recql4_g8_3ng_right3 | 7.256 | 4.893 | 0.356 | 0.252 | 0.674 | 0.708 |
| Recql4_g8_3ng_right4 | 7.972 | 5.59 | 0.338 | 0.155 | 0.701 | 0.459 |
| Recql4_g8_3ng_right5 | 5.666 | 4.363 | 0.239 | 0.141 | 0.770 | 0.590 |
| Recql4_g8_3ng_right6 | 6.550 | 5.398 | 0.325 | 0.248 | 0.824 | 0.763 |
| Recql4_g8_3ng_right7 | 4.637 | 3.399 | 0.245 | 0.105 | 0.733 | 0.429 |
| Recql4_g8_3ng_right8 | 7.136 | 4.509 | 0.315 | 0.277 | 0.632 | 0.879 |
| Recql4_g8_3ng_right9 | 6.141 | 3.89 | 0.293 | 0.109 | 0.633 | 0.372 |
| Recql4_g8_3ng_right10 | 7.016 | 4.533 | 0.352 | 0.137 | 0.646 | 0.389 |
| Recql4_g8_3ng_right11 | 7.165 | 7.132 | 0.341 | 0.32 | 0.995 | 0.938 |
| Recql4_g8_3ng_right12 | 4.700 | 3.933 | 0.238 | 0.218 | 0.837 | 0.916 |
| Recql4_g10_right1 | 4.323 | 3.055 | 0.199 | 0.076 | 0.707 | 0.382 |
| Recql4_g10_right2 | 3.072 | 2.656 | 0.151 | 0.131 | 0.865 | 0.868 |
| Recql4_g10_right3 | 4.032 | 3.294 | 0.180 | 0.139 | 0.817 | 0.772 |
| Recql4_g10_right4 | 3.760 | 3.291 | 0.173 | 0.157 | 0.875 | 0.908 |
| Recql4_g10_right5 | 4.181 | 3.612 | 0.168 | 0.169 | 0.864 | 1.006 |
| Recql4_g10_right6 | 3.293 | 2.739 | 0.182 | 0.135 | 0.832 | 0.742 |
| Recql4_g10_right7 | 3.888 | 3.197 | 0.182 | 0.143 | 0.822 | 0.786 |
| Recql4_g10_right8 | 1.516 | 1.466 | 0.090 | 0.071 | 0.967 | 0.789 |
| Recql4_g10_right9 | 3.456 | 2.548 | 0.164 | 0.174 | 0.737 | 1.061 |
| Recql4_g10_right10 | 2.466 | 2.284 | 0.143 | 0.114 | 0.926 | 0.797 |
| Recql4_g10_right11 | 3.752 | 2.723 | 0.155 | 0.095 | 0.726 | 0.613 |
| Recql4_g10_right12 | 2.564 | 2.076 | 0.139 | 0.137 | 0.810 | 0.986 |
| Recql4_g10_right13 | 2.677 | 2.287 | 0.144 | 0.125 | 0.854 | 0.868 |
| Recql4_g10_right14 | 2.827 | 2.766 | 0.148 | 0.099 | 0.978 | 0.669 |
| Recql4_g10_right15 | 1.502 | 1.390 | 0.103 | 0.061 | 0.925 | 0.592 |
| Recql4_g10_right16 | 4.106 | 3.305 | 0.181 | 0.159 | 0.805 | 0.878 |
| Recql4_g10_right17 | 3.350 | 2.997 | 0.144 | 0.092 | 0.895 | 0.639 |
| Recql4_g10_right18 | 3.692 | 2.969 | 0.178 | 0.108 | 0.804 | 0.607 |
| Recql4_g10_right19 | 4.307 | 3.503 | 0.190 | 0.172 | 0.813 | 0.905 |
| Recql4_g10_right20 | 4.939 | 3.775 | 0.217 | 0.199 | 0.764 | 0.917 |
| Recql4_g10_right21 | 2.744 | 2.159 | 0.143 | 0.135 | 0.787 | 0.944 |
| Recql4_g10_right22 | 1.229 | 1.306 | 0.079 | 0.065 | 1.063 | 0.823 |
| Recql4_g10_right23 | 1.376 | 1.178 | 0.091 | 0.070 | 0.856 | 0.769 |

**Table S3: CRISPant head area proportions descriptive statistics (injected/uninjected side)**

|  | **Control** | ***Recql4* CRISPR /Cas9** | | |
| --- | --- | --- | --- | --- |
|  |  | **2 ng g8** | **3 ng g8** | **2 ng g10** |
| **Sample size N** | 28 | 21 | 24 | 39 |
| **Minimum** | 0.8710 | 0.3720 | 0.6310 | 0.6060 |
| **Maximum** | 1.232 | 0.9120 | 1.096 | 1.063 |
| **Range** | 0.3610 | 0.5400 | 0.4650 | 0.4570 |
| **Mean** | 1.039 | 0.7013 | 0.7605 | 0.8179 |
| **Std. Deviation** | 0.1078 | 0.1157 | 0.1208 | 0.1037 |
| **Std. Error of Mean** | 0.02037 | 0.02525 | 0.02465 | 0.01660 |

**Table S4: Kruskal-Wallis comparison of mean head area proportions (injected/uninjected side)**

| **Dunn's multiple comparisons test** | **Mean rank diff.** | **Significant?** | **Summary** | **Adjusted P Value** |
| --- | --- | --- | --- | --- |
| **Control  vs. 2 ng g8** | 64.98 | Yes | **** | <0.0001 |
| **Control  vs. 3 ng g8** | 55.90 | Yes | **** | <0.0001 |
| **Control  vs. 2 ng g10** | 41.18 | Yes | **** | <0.0001 |

**Table S5:** **CRISPant eye area proportions descriptive statistics (injected/uninjected side)**

|  | **Control** | ***Recql4* CRISPR /Cas9** | | |
| --- | --- | --- | --- | --- |
|  |  | **2 ng g8** | **3 ng g8** | **2 ng g10** |
| **Sample size N** | 28 | 20 | 24 | 39 |
| **Minimum** | 0.9030 | 0.2660 | 0.3720 | 0.3410 |
| **Maximum** | 1.161 | 1.055 | 1.228 | 1.061 |
| **Range** | 0.2580 | 0.7890 | 0.8560 | 0.7200 |
| **Mean** | 1.048 | 0.7443 | 0.7713 | 0.7129 |
| **Std. Deviation** | 0.06583 | 0.2625 | 0.2225 | 0.2003 |
| **Std. Error of Mean** | 0.01244 | 0.05869 | 0.04542 | 0.03208 |

**Table S6: Kruskal-Wallis comparison of mean eye area proportions (injected/uninjected side)**

| **Dunn's multiple comparisons test** | **Mean rank diff.** | **Significant?** | **Summary** | **Adjusted P Value** |
| --- | --- | --- | --- | --- |
| **Control  vs. 2 ng g8** | 44.08 | Yes | **** | <0.0001 |
| **Control  vs. 3 ng g8** | 45.18 | Yes | **** | <0.0001 |
| **Control  vs. 2 ng g10** | 52.56 | Yes | **** | <0.0001 |

**Table S7: Limb phenotype scores from skeletal preparations (Figure 3C)**

| **Group** | **N** | **Normal** | | | | **Reduced** | | | | **Missing** | | | | **Ectopic** | | | |
| --- | --- | --- | --- | --- | --- | --- | --- | --- | --- | --- | --- | --- | --- | --- | --- | --- | --- |
|  |  | LFL | RFL | LHL | RHL | LFL | RFL | LHL | RHL | LFL | RFL | LHL | RHL | LFL | RFL | LHL | RHL |
| **GFP-L** | 14 | 14 | 14 | 14 | 14 | 0 | 0 | 0 | 0 | 0 | 0 | 0 | 0 | 0 | 0 | 0 | 0 |
| **GFP-R** | 20 | 20 | 20 | 20 | 20 | 0 | 0 | 0 | 0 | 0 | 0 | 0 | 0 | 0 | 0 | 0 | 0 |
| **G8 2ng L** | 10 | 1 | 9 | 3 | 10 | 0 | 0 | 1 | 0 | 9 | 1 | 5 | 0 | 0 | 0 | 0 | 0 |
| **G8 2ng R** | 8 | 8 | 2 | 7 | 4 | 0 | 0 | 2 | 0 | 0 | 6 | 0 | 4 | 0 | 0 | 0 | 0 |
| **G8 3ng L** | 24 | 16 | 24 | 20 | 24 | 0 | 0 | 4 | 0 | 7 | 0 | 0 | 0 | 1 | 0 | 0 | 0 |
| **G8 3ng R** | 24 | 24 | 12 | 24 | 20 | 0 | 2 | 0 | 3 | 0 | 8 | 0 | 0 | 0 | 2 | 0 | 1 |

**Table S8: Fore and hindlimb counts of *recql4* sgRNA8 left-sided CRISPants (Figure 6A)**

| ***Recql4* sgRNA8** | **NF stage** | **LFL** | **RFL** | **LHL** | **RHL** |
| --- | --- | --- | --- | --- | --- |
| Left1 | 53 | 0 | 1 | 1 | 1 |
| Left2 | 53 | 0 | 1 | 1 | 1 |
| Left3 | 54 | 0 | 0 | 1 | 1 |
| Left4 | 53 | 0 | 1 | 1 | 1 |
| Left5 | 51 | 0 | 1 | 1 | 1 |
| Left6 | 52 | 1 | 1 | 1 | 1 |
| **Total** | **6** | **1** | **5** | **6** | **6** |

**Table S9: Fore and hindlimb counts of *recql4* sgRNA10 left-sided CRISPants (Figure 6A)**

| ***Recql4* sgRNA10** | **NF stage** | **LFL** | **RFL** | **LHL** | **RHL** |
| --- | --- | --- | --- | --- | --- |
| Left1 | 53 | 1 | 1 | 1 | 1 |
| Left2 | 53 | 0 | 1 | 1 | 1 |
| Left3 | 54 | 0 | 1 | 1 | 1 |
| Left4 | 54 | 0 | 1 | 1 | 1 |
| Left5 | 53 | 0 | 1 | 1 | 1 |
| Left6 | 53 | 0 | 0 | 0 | 1 |
| Left7 | 53 | 0 | 1 | 1 | 1 |
| Left8 | 52 | 0 | 1 | 1 | 1 |
| Left9 | 51 | 1 | 1 | 1 | 1 |
| Left10 | 51 | 0 | 1 | 1 | 1 |
| **Total** | **10** | **2** | **9** | **9** | **10** |

Red text indicates developmental delay of 2 limb bud stages.

**Table S10: Fore and hindlimb counts of *recql4* sgRNA8 right-sided CRISPants (Figure 6A)**

| ***Recql4* sgRNA8** | **NF stage** | **LFL** | **RFL** | **LHL** | **RHL** |
| --- | --- | --- | --- | --- | --- |
| right1 | 53 | 1 | 0 | 1 | 0 |
| right2 | 51 | 1 | 0 | 1 | 0 |
| right3 | 53 | 1 | 0 | 1 | 1 |
| right4 | 51 | 1 | 0 | 1 | 1 |
| right5 | 52 | 1 | 0 | 1 | 0 |
| right6 | 52 | 1 | 0 | 1 | 0 |
| **Total** | **6** | **6** | **0** | **6** | **2** |

**Table S11: Fore and hindlimb counts of *recql4* sgRNA10 right-sided CRISPants (Figure 6A)**

| ***Recql4* sgRNA10** | **NF stage** | **LFL** | **RFL** | **LHL** | **RHL** |
| --- | --- | --- | --- | --- | --- |
| Right1 | 53 | 1 | 1 | 1 | 1 |
| Right2 | 53 | 1 | 0 | 1 | 1 |
| Right3 | 54 | 1 | 0 | 1 | 1 |
| Right4 | 54 | 1 | 0 | 1 | 1 |
| Right5 | 53 | 1 | 0 | 1 | 1 |
| Right6 | 53 | 1 | 0 | 1 | 1 |
| Right7 | 53 | 1 | 0 | 0 | 0 |
| Right8 | 52 | 1 | 0 | 1 | 0 |
| Right9 | 51 | 1 | 0 | 1 | 1 |
| Right10 | 53 | 1 | 0 | 1 | 1 |
| Right11 | 51 | 1 | 0 | 1 | 1 |
| Right12 | 52 | 1 | 0 | 1 | 1 |
| Right13 | 51 | 1 | 1 | 1 | 1 |
| Right14 | 53 | 1 | 0 | 1 | 1 |
| Right15 | 53 | 1 | 0 | 0 | 0 |
| Right16 | 52 | 1 | 0 | 1 | 1 |
| **Total** | **16** | **16** | **2** | **14** | **13** |
