## Supplementary material for "Unilateral loss of *recql4* function in *Xenopus laevis* tadpoles leads to ipsilateral ablation of the forelimb, hypoplastic Meckel’s cartilage and vascular defects": Figure S4

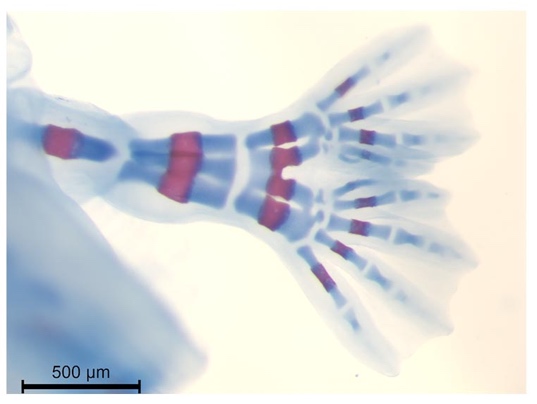

**Figure S4: Ectopic hindlimb skeletal staining. I**ndividual from Figure 4C, stage NF55). There is a single femur, two fibulae and one central tibia, four tarsal bones and two complete hindfeet, with the two copies of digit 1 (hallux) adjacent. Viewed from the dorsal side
