## Supplementary material for "Unilateral loss of *recql4* function in *Xenopus laevis* tadpoles leads to ipsilateral ablation of the forelimb, hypoplastic Meckel’s cartilage and vascular defects": Figure S3

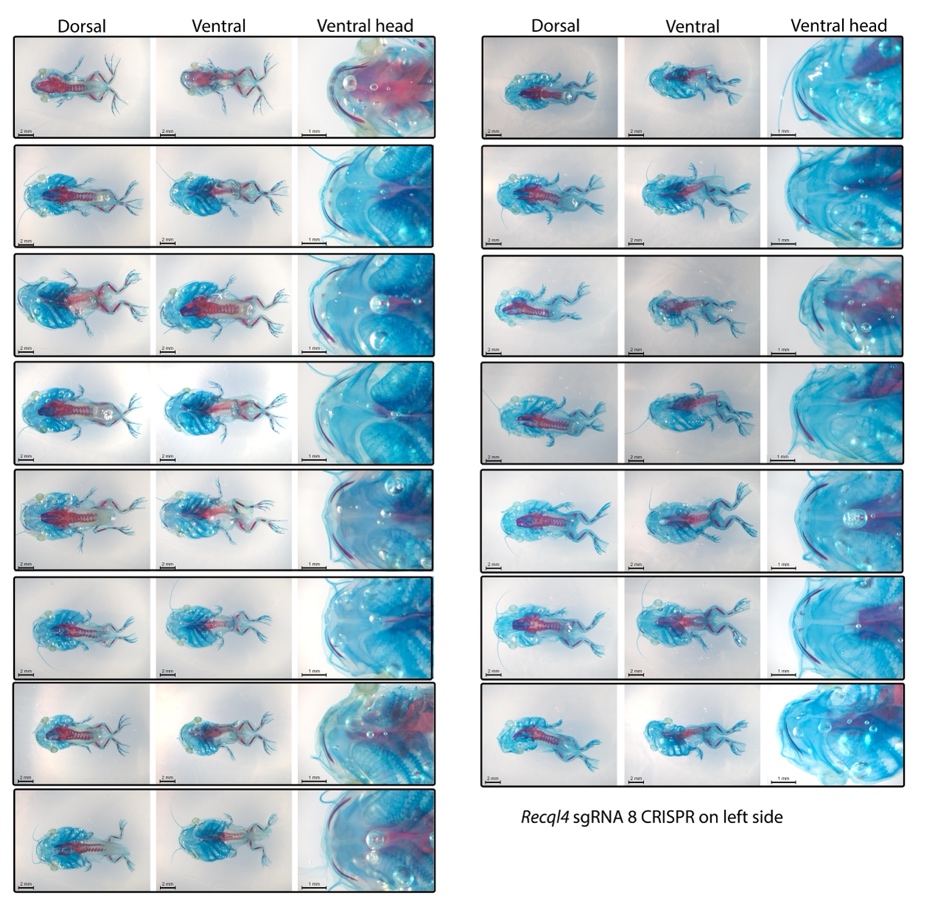

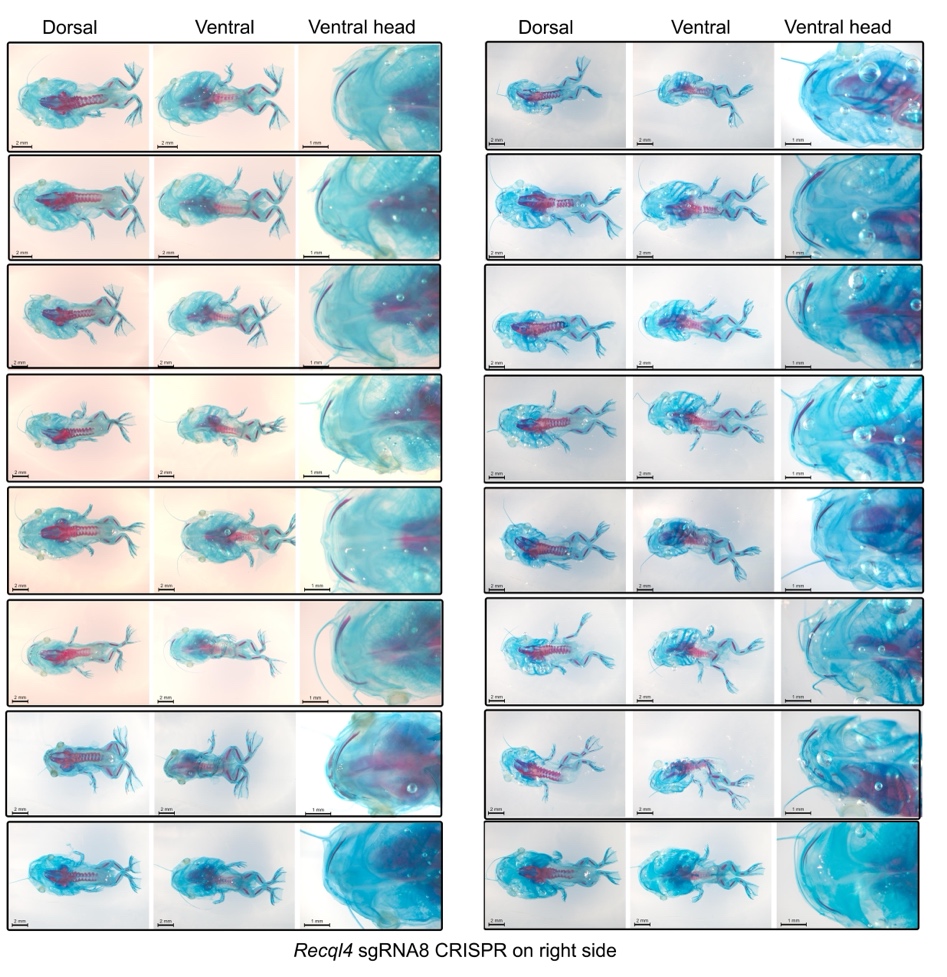

**Figure S3: Cartilage (blue) and bone (red) staining of half recql4 CRISPants (Figures 3, 4)**. Individual tadpoles are boxed, dorsal (left) and ventral (centre) showing whole stage 58 tadpole, right image is a zoom of the head cartilages from the ventral side. Top: left CRISPants, Bottom: right CRISPants.
