## Supplementary material for "Unilateral loss of *recql4* function in *Xenopus laevis* tadpoles leads to ipsilateral ablation of the forelimb, hypoplastic Meckel’s cartilage and vascular defects": Figure S2

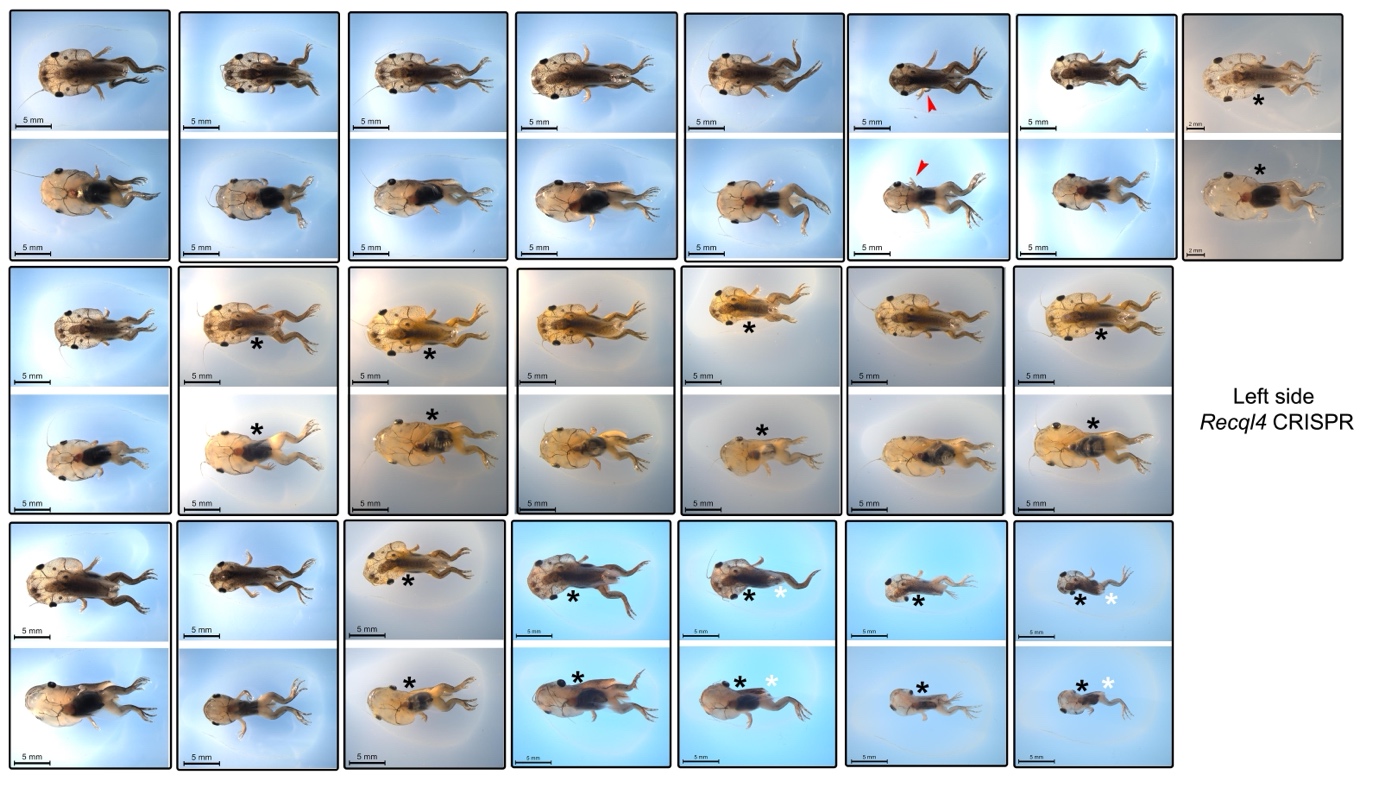

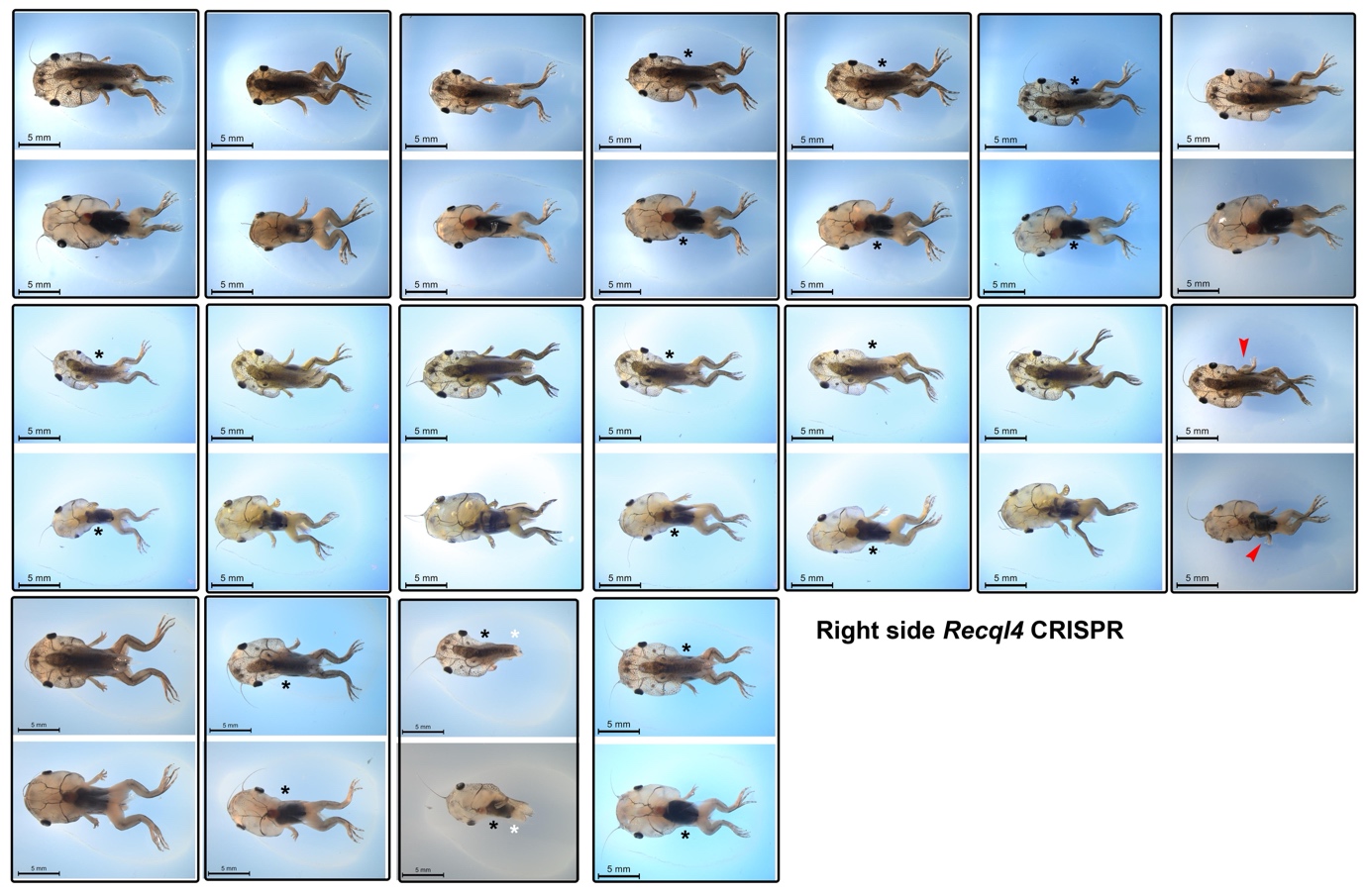

**Figure S2 Stage 58 half CRISPR *recql4* sgRNA8 tadpoles (Figures 3, 4).** Dorsal and ventral views for each animal are boxed, tails have been removed. Top, Left side CRISPants, Bottom, right side CRISPants, red arrowheads indicate ectopic limbs, black asterisk indicates a missing forelimb, white asterisk indicates a missing hindlimb. Scale bars 5 mm.
