## Supplementary material for "Unilateral loss of *recql4* function in *Xenopus laevis* tadpoles leads to ipsilateral ablation of the forelimb, hypoplastic Meckel’s cartilage and vascular defects": Figure S1

| 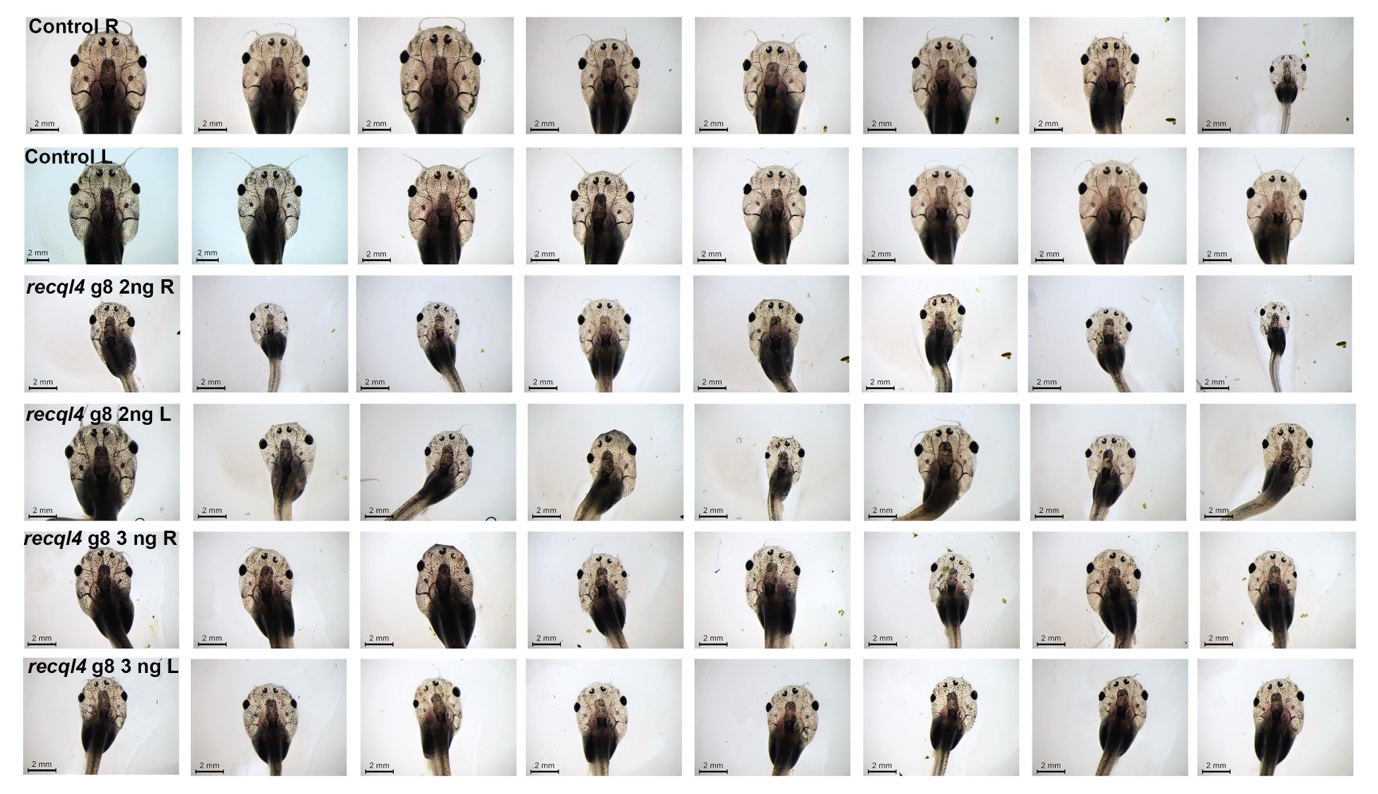 | |
| --- | --- |
| 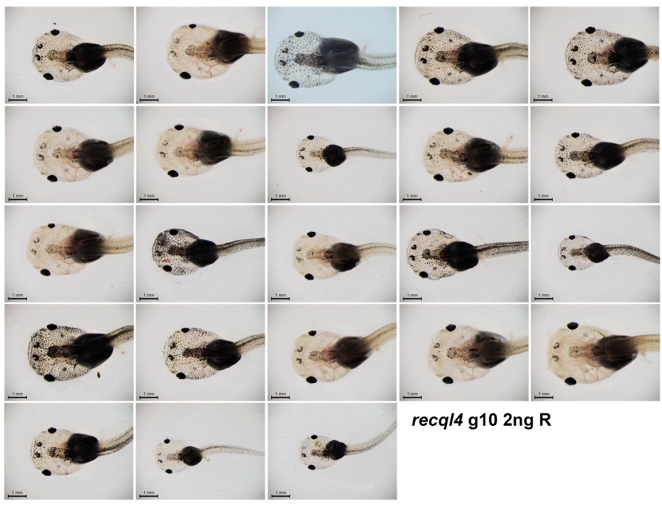 | 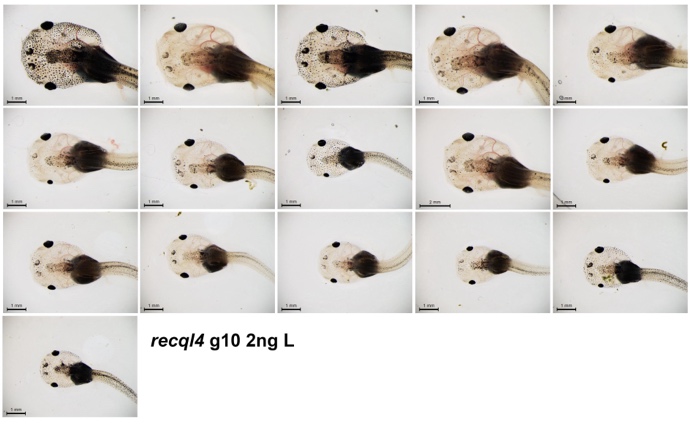 |
| **Figure S1: Examples of *recql4* CRISPR half tadpoles at 3 weeks of development (Figure 2E,F)** Top: first tadpole batch, controls are injected with just GFP mRNA, CRISPants using sgRNA8, scale bars 2 mm. Dorsal view of head and trunk with anterior uppermost. Bottom second tadpole batch using sgRNA10, dorsal view of head and trunk with head to left, scale bars 1 mm. L, injected on Left side, R, injected on right side. | |
